# Exposure assessment suggests some cytotoxic *Bacillus cereus* group genotypes can grow over 3 logs in HTST milk throughout the shelf life at temperature abuse conditions

**DOI:** 10.1101/2024.02.20.581219

**Authors:** Jun Su, Tyler Chandross-Cohen, Chenhao Qian, Laura Carroll, Kayla Kimble, Mackenna Yount, Martin Wiedmann, Jasna Kovac

## Abstract

Cytotoxic *Bacillus cereus* group strains are common causes of foodborne illness including diarrhea. However, our ability to assess food safety risks associated with the exposure to cytotoxic *B. cereus* group strains *via* contaminated food is limited due to the lack of predictive tools. In this study, we experimentally quantified the growth of 17 cytotoxic *B. cereus* group strains, representing six phylogenetic groups, in skim milk broth and used the growth data to develop an exposure assessment model. While none of the tested strains showed detectable growth in HTST milk at 4 or 6°C, 15 of the 17 strains showed growth at 10°C, 1 of the 17 strains showed growth at 8°C, and all strains grew at ≥14°C. Growth data for 16 strains allowed us to generate linear secondary growth models, which were then used to develop the exposure assessment model. We simulated a five-stage supply chain with up to 35 consumer storage days, as that was the timing when distinguishable variations in percent milk containers over 10^5^ CFU/m with different *B. cereus* genotypes were observed. When the initial contamination level of the HTST milk is set at an average of 100 CFU/mL, the model predicts that, on consumer home storage day 21 and 35, 2.81±0.66 and 4.13±2.53 % (mean ± standard deviation) of the milk containers would exceed *B. cereus* group concentrations of 10^5^ CFU/mL; these data represent the average across all strains. Sensitivity analysis showed that variation in the input parameter Q0, the initial physiological state of cells, has the largest effect on model’s prediction for 1 of 4 group II isolates, 1 of 6 group IV isolates and both group V isolates, suggesting the need to better characterize the growth parameters of these isolates. What-if scenario analysis showed that increased mean and variability in storage temperature at the consumer’s home both have substantial influence on final predicted *B. cereus* group concentration in milk containers. This model introduces an initial tool designed to facilitate risk-based food safety decision making for products that are contaminated with low *B. cereus* group levels.

## INTRODUCTION

The *Bacillus cereus* group, also referred to as *B. cereus sensu lato* (*s.l.*), is a complex composed of closely related species that can be categorized into eight phylogenetic groups (Carroll et al., 2020). This group of bacteria raises concerns within the food industry due to their ability to form endospores that withstand heat treatments, such as high-temperature short time (**HTST**) pasteurization of milk (Buss da Silva et al., 2022; Tirloni et al., 2022). Certain strains within this group possess the potential to induce foodborne illness through emetic intoxication, a consequence of the production of heat-stable toxin cereulide within a food matrix. On the other hand, other strains from this group can cause diarrheal intoxication. Upon ingestion by the human host, these bacteria, while growing in the small intestine, produce pore-forming enterotoxins, leading to cell death (Jessberger et al., 2020). Notably, the estimated annual number of foodborne illnesses caused by *B. cereus* group members in the US is reported to be 63,623, with a hospitalization rate of 0.4% (CDC, 2018). Moreover, it is important to acknowledge that foodborne illnesses caused by *B. cereus* group members are often underreported due to their short-lived nature and mild symptoms (Stenfors Arnesen et al., 2008). Most foodborne illnesses caused by the *B. cereus* group have been associated with 10^5^ to 10^8^ cells/spores per gram of food. However, it has been proposed that any food that contains >10^3^ *B. cereus* group cells/spores per gram cannot be considered safe for consumption (EFSA, 2005).

Members of *B. cereus* group are common biological hazards along the dairy production chain, and they often persist in dairy processing environment biofilms (Tirloni et al., 2022). These hazards are particularly relevant to perishable HTST milk, which requires refrigeration along the entire supply chain to ensure product quality and safety. Despite this requirement, HTST milk can be subject to temperature abuses due to an inefficient cold chain. For instance, although efficient temperature control of food products inside refrigerated trucks can usually be achieved, increases in temperature, sometimes to >10°C, during ground operations at the beginning and end of transportation have been frequently reported (Ndraha et al., 2018). Significant temperature abuses have also been observed during consumer transportation from the retail establishment to the home, and inappropriate home storage of food products, which elevate the food safety risk (Mercier et al., 2017). It has been reported that many home refrigerators worldwide are running at higher than recommended temperatures, and this inadequate refrigeration is frequently cited as a factor in incidents of food poisoning (James, Onarinde, and James, 2017). In recent (2003 to 2016) foodborne outbreaks caused by *B. cereus* group strains, most cases were attributed to temperature abuse of food, such as refrigeration temperatures as high as 14°C or long-term storage of food at room temperature (Choi and Kim, 2020).

As *B. cereus* group strains can enter the supply chain at multiple entry points and survive heat treatment, occurrence of low levels of this microorganism in pasteurized milk is not unexpected (Berthold-Pluta et al., 2019; Bartoszewicz, Hansen, and Swiecicka, 2008; Svensson et al., 2004). While the number of *B. cereus* group infections linked to milk and dairy product consumption is unknown, *B. cereus* group-related outbreaks and recalls have been reported. In July 2010, a *B. cereus* linked diarrhea outbreak occurred among workers in a Korean company, with an attack rate of 20.3%. The contamination source was found to be underground water (Choi et al., 2011). Additionally, in 2018, Australian Capital Territory Department of Health had investigated a *B. cereus* related outbreak associated with a multi-course dinner after four people reported gastrointestinal illnesses to a restaurant in Canberra. The beef retained by the restaurant for microbiological analysis showed an unsatisfactory level of *B. cereus* (i.e., a count of 19,000 CFU/g) (Thirkell et al., 2019). Therefore, to facilitate better food safety management decisions about mitigation strategies for the *B. cereus* group, industry needs an exposure assessment tool to quantitatively estimate the dose of *B. cereus* group strains at the point of human consumption, given that a food product is contaminated with a specific concentration and *B. cereus* group genotype (Notermans et al., 1997).

In this project, we modeled the growth of selected cytotoxic *B. cereus* group strains from different phylogenetic groups in contaminated HTST milk along a simulated supply chain and used the data to predict the *B. cereus* group concentrations in a product upon human consumption. We hypothesized that different *B. cereus* group strains have different growth capabilities in HTST milk. Therefore, the percentage of HTST milk containers that contain *B. cereus* group concentrations above a given threshold (i.e., 10^3^ and 10^5^ CFU/mL) on a given storage day differs by the *B. cereus* group strain that contaminates the product. While a more realistic HTST milk shelf life is usually 18-21 days, we modeled a five-stage supply chain with up to 35 consumer storage days, as that was the timing when noticeable variations in percent milk containers over 10^5^ CFU/ml with different *B. cereus* genotypes were observed.

## MATERIALS AND METHODS

### Selection of Bacillus cereus Group Isolates

A total of 17 *B. cereus* group strains were included in this study spanning 6 of the 8 *B. cereus* phylogenetic groups (Carroll et al., 2020). Isolates were selected from a collection of over 300 *B. cereus* group isolates with available genome assemblies to represent (i) diverse virulence gene clusters (i.e., considering the presence and absence of *ces, nhe, hbl, cytk-1, cytk-2, sph, cap, has, bps*), and (ii) phylogenetic groups, which were assigned using BTyper3 v3.3.3 (Carroll et al., 2020). Specifically, BTyper3 was used to calculate average nucleotide identity (ANI) values relative to the species type strain genomes of all validly published and effective *B. cereus* group species published at the time (*n* = 28, accessed March 20, 2023). Using 95 ANI (i.e., the threshold typically used to delineate bacterial species) (Jain et al., 2018), the 17 *B. cereus* group strains used in this study most closely resembled the species type strain genomes of *B. pseudomycoides* (group I isolates PS00125 and PS00135), *albus* (group II isolate PS00193), *tropicus* (group II isolate PS00457), *mobilis* (group II isolate PS00518), *pacificus* (group III isolate PS00474), *cereus* (group IV isolates PS00402, PS00407, PS00433 and PS00495), *thuringiensis* (group IV isolates PS00413 and PS00649), *toyonensis* (group V isolates PS00570 and PS00638) and *cytotoxicus* (group VII isolates PS00194 and PS00536). One strain (PS00564) did not share >95 ANI with any *B. cereus* group species type strain genome; however, using *panC* group assignment, PS00564 belonged to group II (i.e., the phylogenetic group that contains *B. tropicus*), shared >94.0 ANI with the *B. tropicus* type strain genome, and was thus grouped with isolate PS00457 for this study. These selected isolates were grouped into 11 clusters containing unique virulence gene profiles, and further sub-clustered by phylogenetic group. Consequently, we generated 10 sub-clusters that contained cytotoxic isolates (61 cytotoxic isolates in total). Up to two cytotoxic isolates from each virulence-phylogenetic subcluster were selected for inclusion in this study by using a random number generator (Table 1).

**Table 1.** Number of all *B. cereus* group isolates (A) and cytotoxic *B. cereus* group isolates (B) in all clusters and sub-clusters.

| (A) Number of all *B. cereus* group isolates |  |  |  |  |  |  |  |  |  |
| --- | --- | --- | --- | --- | --- | --- | --- | --- | --- |
| Cluster<sup>1</sup> | Phylogenetic group<sup>2</sup> |  |  |  |  |  |  |  | Total |
|  | I | II | III | IV | V | VI | VII | VIII |  |
| 1 | 0 | 0 | 0 | 0 | 0 | 0 | 8 | 0 | 8 |
| 2 | 0 | 0 | 0 | 0 | 1 | 0 | 0 | 0 | 1 |
| 3 | 0 | 5 | 19 | 1 | 0 | 0 | 0 | 0 | 25 |
| 4 | 0 | 0 | 0 | 1 | 0 | 0 | 0 | 0 | 1 |
| 5 | 0 | 5 | 1 | 74 | 3 | 0 | 0 | 0 | 83 |
| 6 | 0 | 1 | 0 | 2 | 0 | 0 | 0 | 0 | 3 |
| 7 | 0 | 0 | 20 | 0 | 0 | 0 | 0 | 0 | 20 |
| 8 | 0 | 16 | 21 | 0 | 0 | 0 | 0 | 0 | 37 |
| 9 | 0 | 19 | 3 | 4 | 36 | 36 | 0 | 4 | 103 |
| 10 | 5 | 0 | 0 | 0 | 1 | 3 | 0 | 0 | 9 |
| 11 | 0 | 0 | 0 | 0 | 1 | 0 | 0 | 0 | 1 |
| (B) Number of cytotoxic *B. cereus* group isolates only |  |  |  |  |  |  |  |  |  |
| Cluster | Phylogenetic group |  |  |  |  |  |  |  | Total |
|  | I | II | III | IV | V | VI | VII | VIII |  |
| 1 | 0 | 0 | 0 | 0 | 0 | 0 | 1 | 0 | 1 |
| 2 | 0 | 0 | 0 | 0 | 0 | 0 | 0 | 0 | 0 |
| 3 | 0 | 3 | 1 | 0 | 0 | 0 | 0 | 0 | 4 |
| 4 | 0 | 0 | 0 | 0 | 0 | 0 | 0 | 0 | 0 |
| 5 | 0 | 1 | 0 | 42 | 0 | 0 | 0 | 0 | 43 |
| 6 | 0 | 0 | 0 | 2 | 0 | 0 | 0 | 0 | 2 |
| 7 | 0 | 0 | 0 | 0 | 0 | 0 | 0 | 0 | 0 |
| 8 | 0 | 0 | 0 | 0 | 0 | 0 | 0 | 0 | 0 |
| 9 | 0 | 1 | 0 | 2 | 4 | 0 | 0 | 0 | 7 |
| 10 | 4 | 0 | 0 | 0 | 0 | 0 | 0 | 0 | 4 |
| 11 | 0 | 0 | 0 | 0 | 0 | 0 | 0 | 0 | 0 |

<sup>1</sup>Isolates are clustered based on unique virulence gene profiles.
<sup>2</sup>Phylogenetic groups assigned by BTyper3 are used for sub-clustering.

### Selection of Temperatures and Sampling Points

Three temperatures, which represent three industrially relevant scenarios, were selected for the growth experiments, which were conducted with 17 cytotoxic *B. cereus* group strains. These include 4°C, the food refrigeration temperature recommended by the Food and Drug Administration (FDA, 2023), 10°C, the refrigeration abuse temperature, and 22°C, the room temperature. For two strains (PS00125 and PS00135), which showed growth at 22°C but not at 10°C, we also collected growth data at 16°C.

In addition, we also conducted growth/no growth experiments at 6 and 8°C for 15 strains that showed growth at 10°C but not at 4°C to better characterize their growth. Two strains (PS00125 and PS00135) that showed growth at 22°C but not at 10°C were also tested at 14 and 18°C. The sampling points for growth experiments at each temperature were selected based on preliminary growth experiments in SMB, which informed the selection of sampling time points to ensure that at least 3 data points were obtained for each growth phase (i.e., lag, exponential growth, and stationary phase).

### Preparation of Spore Suspensions

All strains were streaked from −80°C cryostocks onto brain heart infusion (BHI) agar (BD, cat. No. 241830, Franklin Lakes, NJ), followed by incubating at 30°C for 24 h. A single isolated colony was then inoculated in 5 ml of BHI broth (BD, cat. no. 237500, Franklin Lakes, NJ) in a 15 ml conical tube (VWR, cat. no. 89039-670, Radnor, PA). Inoculated broths were incubated at 30°C for 24 h without shaking. Two separate AK Agar #2 plates (Sporulating Agar) (BD, cat. No. 210912, Franklin Lakes, NJ) were spread-plated with 100 µl of overnight culture and incubated at 30°C for 5 days. To confirm the growth on the AK #2 agar, a wet mount on a microscopy slide was prepared and examined using a phase contrast microscope. After the presence of spores was confirmed, the plates were flooded with 8 ml of chilled sterile deionized water and biomass was scraped with an L-shaped spreader to suspend the spores. The spores were then transferred into new 15 ml conical tubes, and the suspensions were centrifuged (Avanti Centrifuge J-26 XPI) at 10,000 g for 15 minutes. The supernatant was removed, and the spores were resuspended in 3.5 ml of sterile deionized water. Centrifugation was repeated two more times for a total of three washes. After the final wash, the water was removed, and the spores were resuspended in 5 ml of 50% ethanol and incubated at 4°C in a tube rotator (VWR Mini Tube Rotator) running at 15 rpm for 12 h at room temperature to inactivate remaining vegetative cells. After this incubation, the suspension was centrifuged at 10,000 g for 15 minutes. The supernatant was removed, and the spores were resuspended in 3.5 ml of sterile deionized water. Centrifugation was repeated an additional two times for three total washes. After the last centrifugation, the spores were resuspended in 3 ml of sterile deionized water and stored at 4°C for up to 6 months.

To determine the spore suspension concentrations, 100 µl of each spore suspension was transferred to a 1.5 ml microcentrifuge tube (VWR, cat. no. 89000-028, Radnor, PA) and incubated in a water bath at 80°C for 12 minutes, followed by cooling on ice. Using the 1X phosphate buffer saline (PBS), serial dilutions of the spore suspensions were prepared and spread plated, in duplicate, onto BHI agar, followed by a 24 h incubation at 30°C. Colonies on each plate were manually counted, and the CFU/mL of the heat-treated spore suspension was calculated.

### Growth Experiment in Skim Milk Broth

Growth experiment was initiated by preparing a spore suspension at a concentration of 10^3^ CFU/mL in 40 ml of skim milk broth (SMB) (BD, cat. No. 232100, Franklin Lakes, NJ), followed by heat treatment at 80°C for 12 minutes. The growth experiments were conducted in two independent biological replicates, each with two technical replicates, for 17 selected *B. cereus* group strains at 4°C, 10°C and 22°C. At 4°C, samples were collected at time points 0, 96, 144, 192, 240, 288, 384, and 576 h. At 10°C, samples were collected at time points 0, 6, 24, 48, 96, 192, 240, 288, 384, 504 h. At 22°C, data was collected in three experiments with complementary sampling time points, as well as overlapping time points to ensure reproducibility. Specifically, in one experiment, samples were collected at time points 0, 1, 2, 4, 6, 8, and 24 h. In the second experiment, samples were collected at time points 0, 14, 15, 16, 18, 20, and 22 h. In the third experiment, samples were collected at time points 0, 24, 27, and 30 h, with a 32 h time point being added for isolates that had not reached the stationary phase at 30 h. Growth experiments conducted at 16°C for two strains (PS00125 and PS00135) were also conducted in two biological replicates, each with two technical replicates, with data collected at time points 0, 1, 2, 4, 6, 8, 14, 18, 24, 48, and 96 h.

At each sampling time point, 1 ml of the sample was serially 10-fold diluted using a 1X PBS buffer and 100 μl of dilutions 10^-1^, 10^-3^, & 10^-5^ were plated on BHI agar plates, in duplicates. The BHI agar plates were incubated at 30°C for 12 to 24 h before colony enumeration. Isolates with filamentous colony morphology were pour-plated to prevent colony spreading and were incubated at 30°C for 12 to 24 h. The colonies on countable plates, with 30 to 300 colonies, were manually counted. Plating of 100 μl SMB and 100 μl 1X PBS were included in each experiment as negative controls. Experiments were not considered valid if the negative controls had observable colony growth.

### Growth/No Growth Experiments

For growth/no growth experiments, spore suspensions were inoculated in SMB at a concentration of 10^3^ CFU/mL, and incubated at 6°C or 8°C for 504 h. The data was collected at time points 0 and 504 h. Experiments were conducted with two independent biological replicates, each with two technical replicates. For two strains that showed growth at 22°C but not at 10°C, the growth/no growth experiments were also conducted at 14°C and 18°C. Spore suspensions of these strains were inoculated in SMB at a concentration of 10^3^ CFU/mL, and incubated for 384 h. The data was collected at time points 0 and 384 h. Experiments were conducted with two independent biological replicates, each with two technical replicates. Microbial concentration was determined as described in the previous section. An increase of cell concentration by 1 log CFU/mL during the growth/no growth experiment was considered growth.

### Primary Model Fitting

To estimate the growth parameters, a Baranyi model (Baranyi and Roberts, 1994) was fitted to growth data for 17 selected *B. cereus* group strains collected at 22°C, 16°C and 10°C using the packages “nlsMicrobio” version 0.0-3 (Baty, 2022) and “minpack.lm” version 1.2-3 (Elzhov, 2022) in R version 4.1.2 (R Core Team, 2021). The Baranyi model was selected as the primary model since it was the one used for microbial growth prediction under dynamic environmental conditions in R package “biogrowth” version 1.0.2 (Garre, 2022). This primary model was used for exposure assessment model development to facilitate prediction of growth under dynamic temperature conditions. The equation (Eq. 1) is presented below:

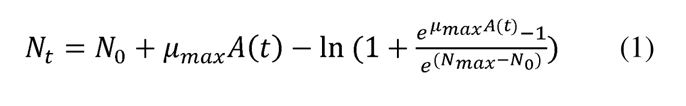

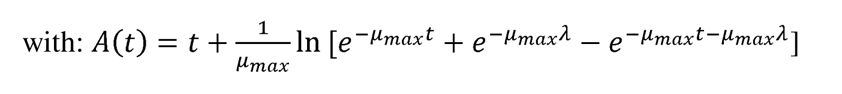

where µ_max_ is the maximum specific growth rate in ln/h, λ is lag time in h, N_0_ is the initial population in log_10_ CFU/mL and N_max_ is the maximum population reached in log_10_ CFU/mL. The last three data points of isolate PS00413 replicate 1 at 10°C were excluded when fitting Baranyi model since cells had entered the death phase (Supplemental Material Figure S1).

### Secondary Model Fitting

A reduced Ratkowsky model was selected as the secondary model to calculate T_min_ and b (Ratkowsky et al., 1982), since our available data was confined to growth under sub-optimal temperatures. The unit of µ_max_ was converted to log_10_/day before fitting the secondary model in R version 4.1.2 (R Core Team, 2021). The equation (Eq. 2) is presented below:

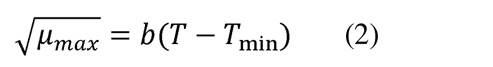

where T is temperature represented in °C, b is the slope and T_min_ is the theoretical minimum growth temperature represented in °C.

Growth/no growth experiment data at 6°C, 8°C, and 10°C were also used as input data to construct the secondary model. Specifically, for strains that showed no growth at 6°C, 8°C, or 10°C, µ_max_ was assumed to be 0 at the given temperature and this µ_max_ value was used as an input when fitting the secondary growth model.

### Statistical Analysis

Kruskal-Wallis tests were performed on the λ, μ_max_ and N_max_ of all strains whose growth data were obtained from both 22°C and 10°C growth experiments. A Bonferroni-corrected Dunn’s test was used to identify the pairs of strains that showed significant differences in lag time and maximum growth rate. The non-parametric tests were applied because the growth parameters for these strains failed to meet normality and equal variance assumptions.

### Exposure Assessment Model Overview

The exposure assessment model simulates the growth of selected cytotoxic *B. cereus* group strains in one lot of HTST milk transported along a five-stage supply chain that includes (i) processing facility storage, (ii) transportation from the processing facility to retail, (iii) retail storage, (iv) transportation from retail to the consumer’s home, and (v) consumer storage for up to 35 days. Each hypothetical lot of HTST milk was contaminated by one *B. cereus* group strain and had 100 half-gallon milk containers represented by 100 iterations. This model predicts *B. cereus* group concentrations in milk containers on days 14, 21, and 35 of consumer storage. Model outputs were also used to compute the percentage of milk containers with *B. cereus* group concentrations that exceeded 10^3^ and 10^5^ CFU/mL at days 14, 21, and 35.

### Exposure Assessment Model Parameters

The input parameters for the exposure assessment model include initial contamination levels, growth parameters of selected *B. cereus* group strains, and the temperature profiles for each stage of the supply chain. The initial contamination levels (N_0_) for 100 iterations were generated from a Poisson distribution in R version 4.1.2 (R Core Team, 2021) with a lambda (i.e., mean) value of 100 CFU/mL. The seed for the initial contamination level generation was set to 42, for consistency.

The initial physiological state of cells (Q0) was calculated as shown in Eq. 3 and Eq. 4 and described in the Baranyi model (Baranyi and Roberts, 1994):

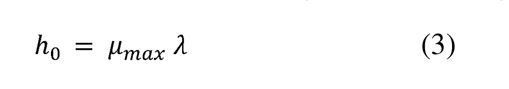

where µ_max_ is the specific maximum growth rate in ln/h, λ is the lag time in h.

h_0_ was averaged across 2 replicates at 22°C and 2 replicates at 10°C for each strain to obtain h_av_ which was used to calculate Q0 specific to each strain:

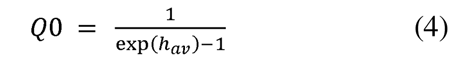

The maximum microbial population (N_max_) for each given *B. cereus* group strain was estimated from the Baranyi model and averaged across all replicates from growth experiments at different temperatures for a given *B. cereus* group strain.

The maximum growth rate at the optimum growth temperature (µ_opt_) was calculated as shown in Eq. 5:

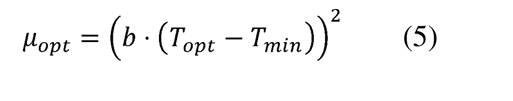

where T_min_ and b were calculated from the secondary model. T_opt_ is the optimum growth temperatures by clade previously reported (Nakamura, 1998, Baranyi et al., 2017). The unit of μ_opt_ is in log_10_/day.

A unique time and temperature profile for every milk container in each lot was generated by randomly drawing a time-temperature combination from the corresponding time and temperature distributions (see Table 2) at each stage of the supply chain as described in a previous study (Qian et al., 2023). The seed was set to 1, for consistency.

**Table 2.** Variables used in the time-temperature distributions of the five-stage supply chain.

| Symbol | Variable name | Distribution | Unit | Reference |
| --- | --- | --- | --- | --- |
| Stage 1. Processing plant storage |  |  |  |  |
| $t_F$ | Storage time of milk containers | Uniform (1,2) | d | (Qian et al., 2023) |
| $T_F$ | Storage temperature of milk containers | Uniform (3.5, 4.5) | °C | (Qian et al., 2023) |
| Stage 2. Transportation from processing plant to retail |  |  |  |  |
| $t_T$ | Storage time of milk containers | Triangular (1, 10, 5) | d | (FDA and Health Canada, 2015) |
| $T_T$ | Storage temperature of milk containers | Triangular (1.7, 10.0, 4.4) | °C | (FDA and Health Canada, 2015) |
| Stage 3. Retail storage |  |  |  |  |
| $t_S$ | Storage time of milk containers | TruncNormal (0.042, 10.0, 1.821, 3.3)<sup>1</sup> | d | (Qian et al., 2023) |
| $T_S$ | Storage temperature of milk containers | TruncNormal (-1.4, 5.4, 2.3, 1.8) | °C | (FDA and Health Canada, 2015) |
| Stage 4. Transportation from retail to consumer's home |  |  |  |  |
| $t_{T2}$ | Storage time of milk containers | TruncNormal (0.01, 0.24, 0.04, 0.02) | d | Expert opinion |
| $T_{T2}$ | Storage temperature of milk containers | TruncNormal (0, 10.0, 8.5, 1.0) | °C | Expert opinion |
| Stage 5. Consumer storage |  |  |  |  |
| $t_H$ | Storage time of milk containers | 1-35 | d | |
| $T_H$ | Storage temperature of milk containers | TruncLaplace (-1, 15, 4.06, 2.31)<sup>2</sup> | °C | (Pouillot et al., 2010) |

<sup>1</sup> In a TruncNormal (a, b, c, d) distribution, a = min, b = max, c = mean, and d = standard deviation.
<sup>2</sup> In a TruncLaplace (a, b, c, d) distribution, a = min, b = max, c = location parameter, and d = dispersion.

### Sensitivity Analysis

A sensitivity analysis was performed to assess the impact of input parameters, specifically Q0 and N_max_, on the model’s outcome, which is the percent milk containers exceeding *B. cereus* group concentrations of 5 log on consumer storage day 35. In addition to the average h_0_, both the minimum and maximum h_0_ were employed to determine the corresponding maximum and minimum Q0. The input parameters for running the model on selected strains included the minimum, average, and maximum values for Q0 (representing the physiological state of the cell, calculated as the product of growth rate and lag time). Similarly, the minimum, average, and maximum values for N_max_ were used as input parameters for running the model on the selected strains.

### What-If Scenario

Previous studies (Ndraha et al., 2018) suggest that temperature abuse along the HTST milk supply chain is most likely to occur during product transportation (from processing facility to retail or from retail to the consumer’s home) and during consumer home storage. Here, we simulated four possible scenarios that could occur along an HTST milk supply chain including: (i) a delivery delay by 3 days when HTST milk products were transported from processing facility to retail, with no temperature abuse (e.g., amid a severe snowstorm in New York, milk delivery was delayed by 3 days due to impassable roads, safety concerns, and the time required for road clearing and recovery operations; the products were kept in a well-controlled distribution center during the delay), (ii) a temperature abuse simulating a consumer’s transportation of HTST milk products from retail to home in a hot summer, which results in an average of 33.5°C (i.e., 25°C higher than the average temperature of the base model), (iii) a slight temperature deviation in a consumer’s home refrigerator, modelled as a 1°C temperature increase (i.e., an average home storage temperature of 5.45°C), and (iv) an increase in the variability of refrigeration temperatures due to unstable power supply (e.g., rolling blackout in California during summer), modelled as a 1°C increase in the dispersion parameter of the distribution of consumer home storage temperature. Percent milk containers that had *B. cereus* group concentrations >10^5^ CFU/ml on consumer storage day 21 and 35 were calculated for all four scenarios and compared to the base model prediction (Table 3).

**Table 3.** Implementation of what-if scenarios and outcomes.

| Scenario | Implementation<sup>1</sup> | Modified time/temperature distribution<sup>1</sup> | Percent milk containers exceeding 5 logs on consumer storage day 35 (mean ± standard deviation) |
| --- | --- | --- | --- |
| **Baseline** | N/A | N/A | 4.13±2.53 |
| **Scenario (i)** |  |  |  |
| A three-day delay on HTST milk product delivery from processing facility to retail without temperature abuse | Shift in the distribution of transportation time from processing plant to retail 3 units towards the right | $t_T \sim \text{Triangular}(4, 13, 8)$ | 4.13±2.53 |
| **Scenario (ii)** |  |  |  |
| A temperature abuse when a consumer transported the HTST milk products from retail back home in a hot summer | Shift in the distribution of transportation temperature from retail to consumer's home 25 units towards the right | $T_{T2} \sim \text{TruncNormal}(25, 35, 33.5, 1.0)$ | 4.44±3.03 |
| **Scenario (iii)** |  |  |  |
| A slight temperature deviation in consumer's home refrigerator | Shift in the distribution of consumer home storage temperature 1 unit towards the right | $T_H \sim \text{TruncLaplace}(0, 16, 5.06, 2.31)$ | 8.88±3.84 |
| **Scenario (iv)** |  |  |  |
| An increase in the variability of refrigeration temperatures at consumer's home | Increase in the dispersion parameter of the distribution of consumer home storage temperature by 1 unit | $T_H \sim \text{TruncLaplace}(-1, 15, 4.06, 3.31)$ | 10.81±3.15 |

<sup>1</sup>Details of variables used in the time-temperature distributions of the five-stage supply chain can be found in Table 2.

### Model Programming and Data Availability

The model was developed in R version 4.1.2 (R Core Team, 2021) with the modified “biogrowth” package version 1.0.2 (Garre, 2022). The seed for Monte Carlo simulation was set to 1, for consistency. The raw data used for model construction and R code for the entire study are available at https://github.com/FSL-MQIP/Bacillus-cereus-exposure-assessment-model.

## RESULTS

### Bacillus cereus Group Strains Differ in Growth Characteristics, Including the Observed Minimum Growth Temperature

The 17 selected *B. cereus* group strains (representing six phylogenetic groups) differed in terms of their minimum growth temperature in SMB. While no strain showed detectable growth (i.e., > 1 log increase in CFU/ml) at 4°C, 15 strains showed growth at 10°C; 14 of these strains did not grow at 6 or 8°C, but one strain (PS00638 from group V) grew at 8°C. The two strains that did not show growth at 10°C showed growth at 14 and 16°C, as determined based on the growth/no growth experiments at these two temperatures. Hence, the experimentally obtained growth boundaries for the 17 strains assessed here were (i) 8°C (1 strain), (ii) 10°C (14 strains), and (iii) 14°C (2 strains, which represented the only two group I strains characterized in this study).

Due to different minimum growth temperatures associated with the 17 tested strains, the following datasets were available to estimate growth parameters (i.e., lag time, maximum growth rate, maximum microbial population): (i) 22°C for all 17 strains; (ii) 10°C for 15 strains, and (iii) 16°C for 2 strains. For 22°C, parameter estimates obtained after fitting the growth data with the Baranyi model were (i) lag time ranging from 0 h to 22.04 h, (ii) maximum growth rate ranging from 0.31 ln/h to 2.02 ln/h; and (iii) maximum microbial population ranging from 5.24 log_10_ CFU/mL to 7.61 log_10_ CFU/mL. By comparison, for 10°C, parameter estimates were (i) lag times from 10.02 h to 194.05 h, (ii) maximum growth rates from 0.03 ln/h to 0.34 ln/h and (iii) maximum microbial population from 5.01 log_10_ CFU/mL to 7.37 log_10_ CFU/mL. At 16°C, parameter estimates were (i) lag times from 9.12 h to 14.02 h, (ii) maximum growth rates from 0.35 ln/h to 0.95 ln/h and (iii) maximum microbial populations from 6.17 log_10_ CFU/mL to 6.28 log_10_ CFU/mL (Supplemental Material Table S1).

### B. cereus Phylogenetic Groups Differ in Growth Characteristics, including the Maximum Growth Rate at 22°C and Hypothetical Minimum Growth Temperature

To further evaluate *B. cereus* group growth characteristics, the growth parameters obtained for different strains were analyzed based on phylogenetic groups (i.e., group I, II, IV, V, and VII; see Fig. 1). Group III was excluded from these analyses since it only contained one strain. The Kruskal-Wallis test for 22°C growth parameters indicated that both lag time and maximum growth rate differed significantly among phylogenetic groups (p-values of 0.040 and 0.036, respectively), while maximum microbial population did not differ significantly (p-values > 0.1). Group I strains showed the highest median maximum growth rate (0.92 ln/h), which was significantly (p = 0.024) higher than that for group IV strains (0.48 ln/h). With regard to the lag time, group VII strains had the longest median lag time (20.30 h), whereas group IV strains had the shortest median lag time (8.43 h).

**Figure 1.**
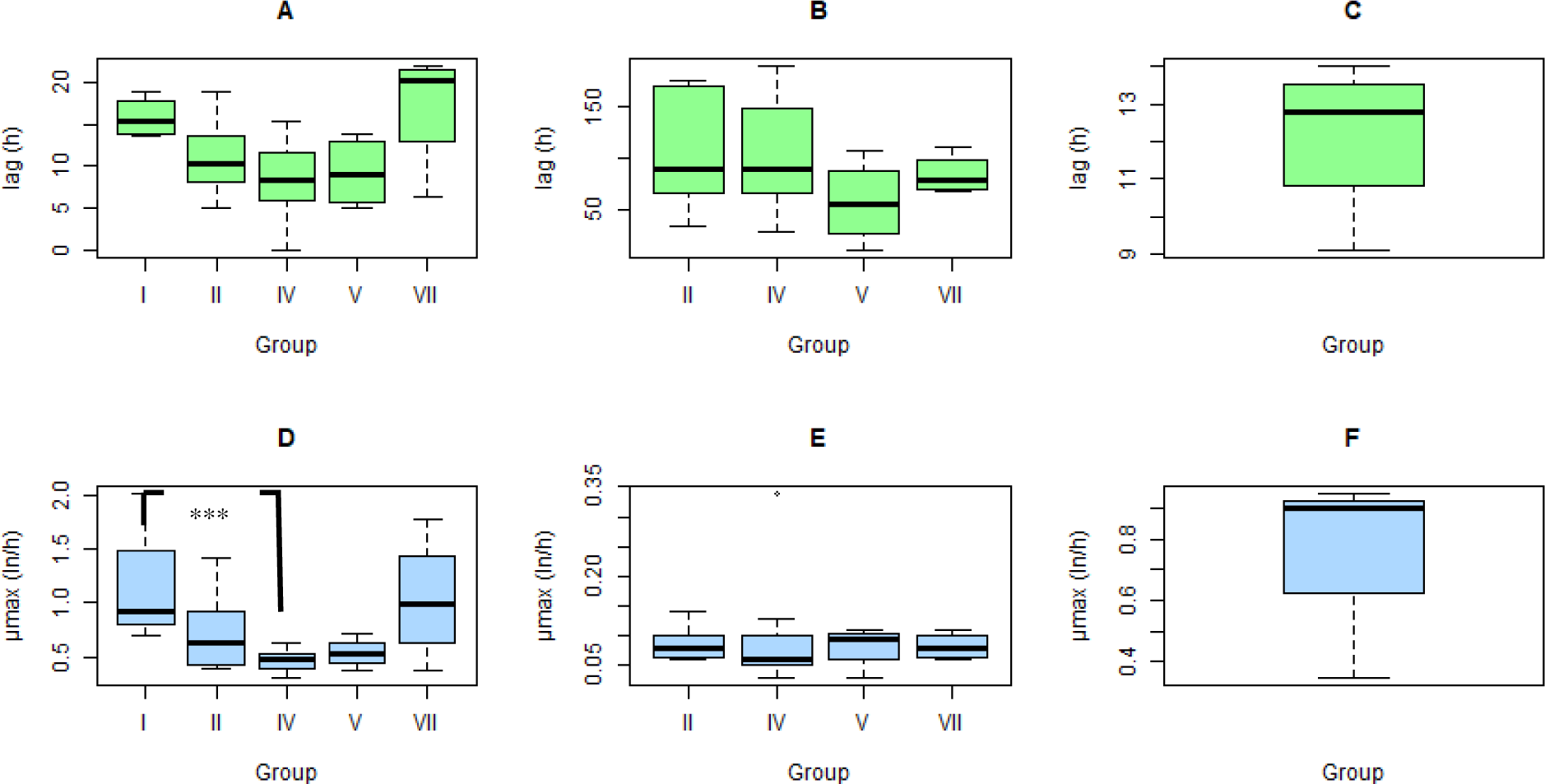
Boxplot that shows the within and between group variability of growth parameters (i.e., lag and µ_max_ at 22°C for (A, D) 16 *B. cereus* group strains representing 5 phylogenetic groups [group I (n=4), group II (n=8), group IV (n=12), group V (n=4), group VII (n=4)], (B, E) 14 *B. cereus* group strains at 10°C, representing 4 phylogenetic groups, and (C, F) 2 *B. cereus* group strains from group I at 16°C (group I strains did not grow at 10°C)). Group III was excluded from this plot since it only had one strain. ***P < 0.05 based on post hoc Dunn’s test performed following a Kruskal-Wallis test that found μ_max_ to be significantly different between group I and group IV strains when grown at 22°C.

By comparison, the Kruskal-Wallis test for 10°C growth parameters showed no significant differences (p-values > 0.1) for any of the three parameters (i.e., lag time, maximum growth rate, maximum microbial population) among phylogenetic groups. Numerically, group V strains had the highest median maximum growth rate (0.10 ln/h), whereas group IV strains had the lowest median maximum growth rate (0.06 ln/h). Group II strains had the longest median lag time of 89.11 h, while group V strains had the shortest median lag time of 53.94 h (Fig. 1). [Figure 1]

To further assess differences in secondary growth parameters between groups, we fitted maximum growth rates for all 17 strains with a secondary growth model (i.e., reduced Ratkowsky model). This model was converged using data from all strains except isolate PS00135 (group I) that had a non-linear relationship between maximum growth rate and temperature under sub-optimal growth conditions (Supplemental Material Fig. S2). We therefore were not able to calculate T_min_ (i.e., hypothetical minimum growth temperature) for this strain. For the rest of the strains, T_min_ ranged from 4.26°C to 9.22°C. The average T_min_ of strains from groups II, III, IV, V and VII were similar, ranging from 5.47°C to 6.98°C. The only strain from group I (PS00125) showed a substantially higher T_min_ (9.22°C) compared to the strains from other groups (Table 4).

**Table 4.**
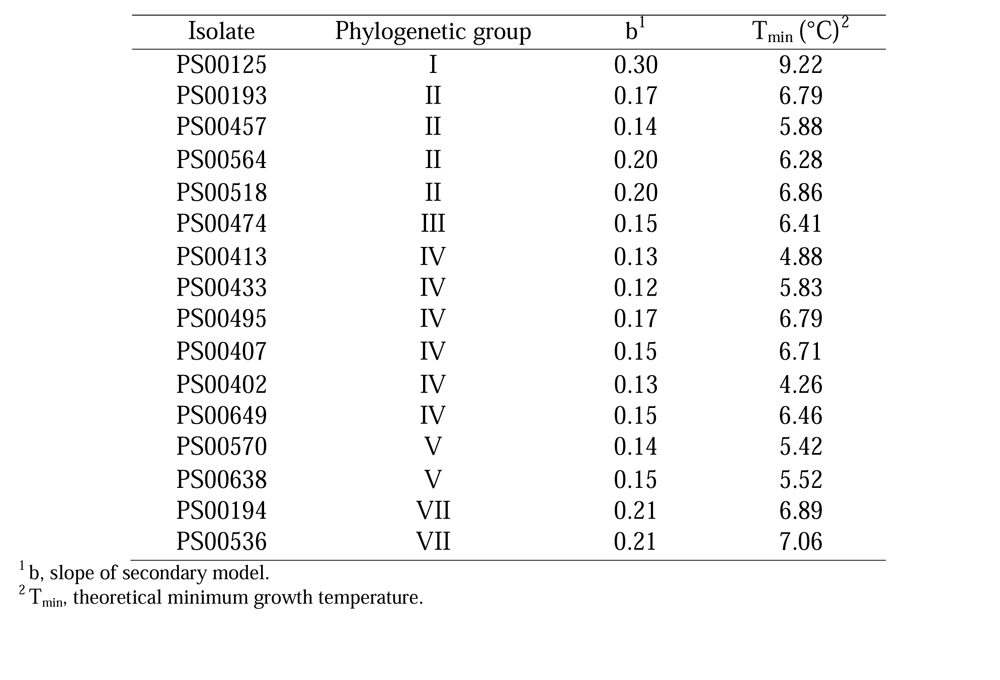
The slope of secondary model and theoretical minimum growth temperature for 16 isolates.

| Isolate | Phylogenetic group | b<sup>1</sup> | T<sub>min</sub> (°C)<sup>2</sup> |
| --- | --- | --- | --- |
| PS00125 | I | 0.30 | 9.22 |
| PS00193 | II | 0.17 | 6.79 |
| PS00457 | II | 0.14 | 5.88 |
| PS00564 | II | 0.20 | 6.28 |
| PS00518 | II | 0.20 | 6.86 |
| PS00474 | III | 0.15 | 6.41 |
| PS00413 | IV | 0.13 | 4.88 |
| PS00433 | IV | 0.12 | 5.83 |
| PS00495 | IV | 0.17 | 6.79 |
| PS00407 | IV | 0.15 | 6.71 |
| PS00402 | IV | 0.13 | 4.26 |
| PS00649 | IV | 0.15 | 6.46 |
| PS00570 | V | 0.14 | 5.42 |
| PS00638 | V | 0.15 | 5.52 |
| PS00194 | VII | 0.21 | 6.89 |
| PS00536 | VII | 0.21 | 7.06 |

<sup>1</sup> b, slope of secondary model.
<sup>2</sup> T<sub>min</sub>, theoretical minimum growth temperature.

### The Base Model Predicts a Higher Percent of Containers Exceeding B. cereus Group Concentrations of 10^5^ and 10^3^ CFU/ml for One Group IV Strain and One Group V Strain

The growth data for the 16 *B. cereus* group strains that showed linear secondary models were used for the development of an exposure assessment model. The base model underwent a “sanity check” to assess the growth predictions for 16 *B. cereus* group strains in HTST milk products stored at 3°C for 90 days, with an initial contamination level of 100 CFU/mL. This temperature was selected because it was about 1°C below the lowest T_min_ of all strains calculated from the secondary model. As expected, the model predicted no growth for any of the 16 strains at 3°C, which aligns with the experimental outcomes that none of the strains had detectable growth at 4°C.

The exposure assessment model was subsequently used to predict growth in HTST milk for each of the 16 included strains up to day 35 of consumer storage, using temperature profiles that include different time and temperature distributions for 5 stages of the supply chain (Table 2). The resulting data were used to compute the percentage of simulated milk containers that had *B. cereus* group concentrations >10^5^ CFU/ml and 10^3^ CFU/ml on consumer home storage day 14, 21, 35, respectively. The percentage of milk containers exceeding 10^5^ CFU/ml on day 14, 21 and 35 were 1.94±0.85 (mean ± standard deviation), 2.81±0.66 and 4.13±2.53, respectively, while the percentage of milk containers exceeding 10^3^ CFU/ml on the same days were 3.00±0.82, 3.88±2.25 and 5.63±3.67.

Model predictions for individual strains (Fig. 2), showed limited differences among strains with regard to percentage of containers containing above 10^5^ CFU/ml (Fig. 2a) at consumer storage days 14 and 21. However, at day 35, four strains yielded >5% of containers that exceeded 10^5^ CFU/ml, including two strains that yielded approximately 10% of containers that exceeded 10^5^ CFU/ml (isolates PS00413 [group IV] and PS00638 [group V]). The same four strains that showed a higher percentage of containers above 10^5^ CFU/ml also showed a higher percentage of containers above 10^3^ CFU/ml (Fig. 2b).

**Figure 2.**
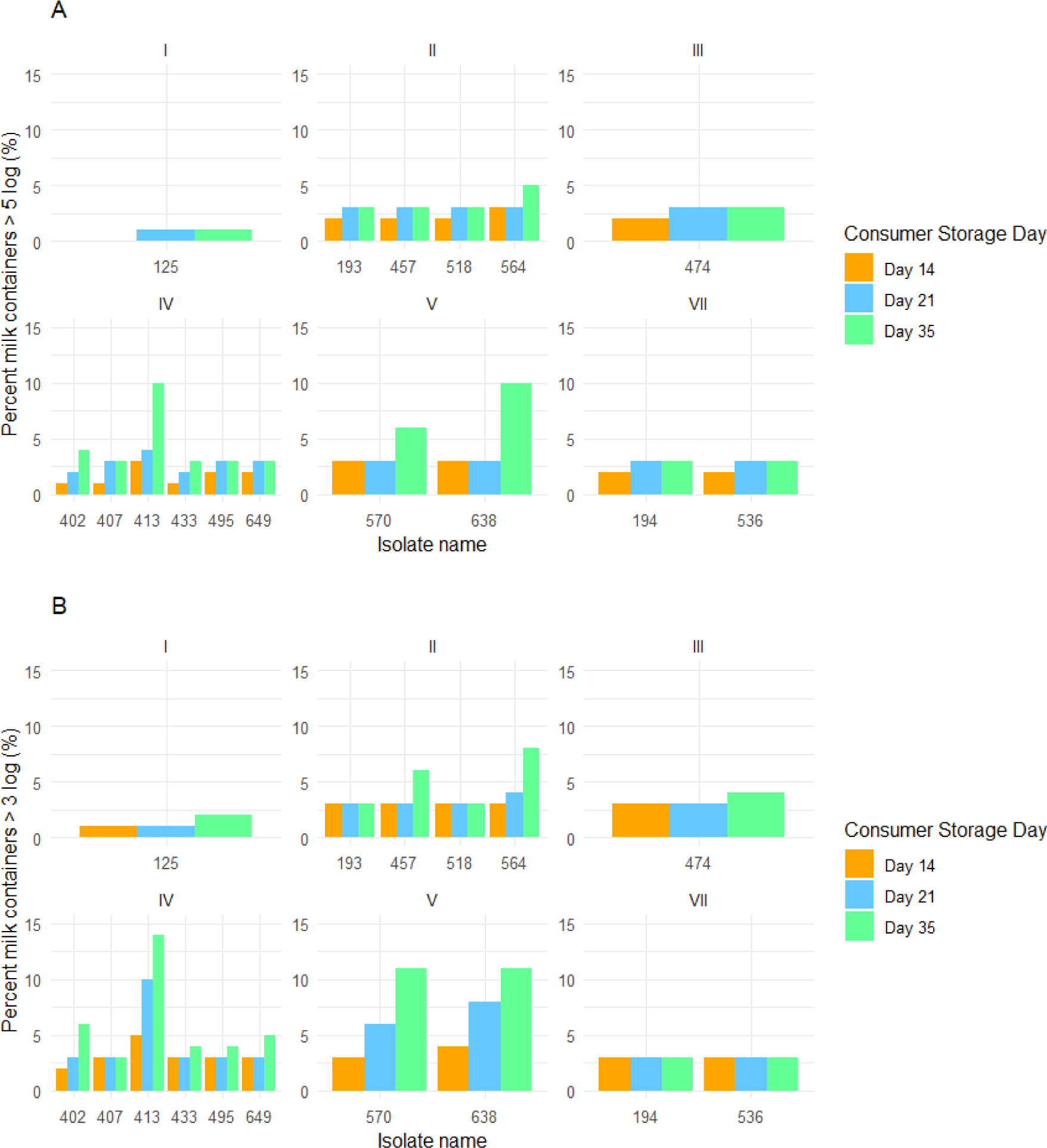
Base model prediction results. The percentage of simulated milk containers contaminated with a given *B. cereus* group (i.e., I, II, III, IV, V, VII) that exceeded 5 log_10_ CFU/ml (A) and 3 log_10_ CFU/ml (B) on consumer home storage day 14, 21, or 35.

Our analyses identified five strains for which the percentage of containers above 10^3^ CFU/ml (i.e., 3%) was the same at consumer storage days 14, 21, and 35 (i.e., isolates PS00193 and PS00518 from group II, isolate PS00407 from group IV, and isolates PS00194 and PS00536 from group VII). These five strains, as well as four additional strains (i.e., isolate PS00457 from group II, isolate PS00474 from group III, isolates PS00495 and PS00649 from group IV) also showed the same percentage (i.e., 3%) of containers above 10^5^ CFU/ml at consumer storage days 21 and 35. We further evaluated the conditions for strains that had the same model predictions at different consumer storage time to verify our model is performing reasonably. An analysis of the 100 model simulations for these nine strains (which represented 3 group II strains, 1 group III strain, 3 group IV strains, and 2 group VII strains) showed that for each of these strains, only 3 out of 100 simulated temperature profiles had a consumer home storage temperature >10°C. For the other 97 simulations, the consumer storage temperature was either (i) below T_min_ of these nine strains (and thus resulted in no predicted growth) or (ii) above T_min_ but below 10°C, a temperature at which our model predicts that these isolates grew so slowly that they would not exceed 10^5^ CFU/ml by consumer storage day 35.

### Sensitivity Analysis Revealed that Variation in the Input Parameter Q0 Significantly Influenced the Model Predictions for Certain Strains

In the sensitivity analysis, we evaluated the impact of both N_max_ and Q0 on the percent milk containers exceeding *B. cereus* group concentrations of 10^5^ CFU/ml on consumer storage day 35 for 16 strains. This combination of consumer storage day and concentration cut-off was used because 10^5^ CFU/ml represents the *B. cereus* regulatory threshold in several countries and day 35 showed some of the largest variations in model predictions between strains. The sensitivity analysis for N_max_ predicted an average of 4.06±2.54 (mean ± standard deviation) and 4.13±2.53% milk containers exceeding *B. cereus* group concentrations of 10^5^ CFU/ml on consumer storage day 35 when using the minimum and maximum N_max_ as the input parameters, respectively, as compared to 4.13±2.53% when using the average N_max_. Only one out of the 16 strains (PS00402; group IV) showed detectable changes in the percent milk containers over 10^5^ CFU/ml. For this strain, the percent milk containers over 5 logs dropped by 1% point when using the minimum N_max_ as the input as compared to using the average N_max_ (Supplement al Material Table S2). Our data thus indicate that model outcomes showed limited sensitivity to changes in N_max_.

In contrast, the model outcomes were more sensitive to the parameter Q0. Using the minimum and maximum Q0 as the input parameters in the sensitivity analysis resulted in an average of 3.00±2.42 (mean ± standard deviation) and 6.31±4.14% milk containers exceeding *B. cereus* group concentrations of 10^5^ CFU/ml on consumer storage day 35, respectively, as compared to 4.13±2.53% when using the average Q0. This analysis also revealed that the sensitivity of model outcomes to variation in Q0 differed substantially by strain. The four strains where model outcomes were most sensitive to changes in Q0 were isolates PS00402 (group IV), PS00638 (group V), PS00564 (group II), and PS00570 (group V). On the other hand, the model outcomes were not sensitive to the change in Q0 for three strains from group IV and the same predictions were generated with varying Q0 values (Fig. 3). This is because (i) one strain (i.e., isolate PS00433 from group IV) had lower variability in the parameter Q0 compared to other strains, and (ii) our model predicted that the other two strains (i.e., isolates PS00407 and PS00495 from group IV) grew very slowly and would not reach >10^5^ CFU/ml on consumer storage day 35 under storage temperature of <10°C even in the worst-case situation (i.e., highest experimentally determined Q0). Overall, the sensitivity analysis indicated that additional growth data collection for strains with great uncertainty in Q0 is necessary to improve the prediction precision.

**Figure 3.**
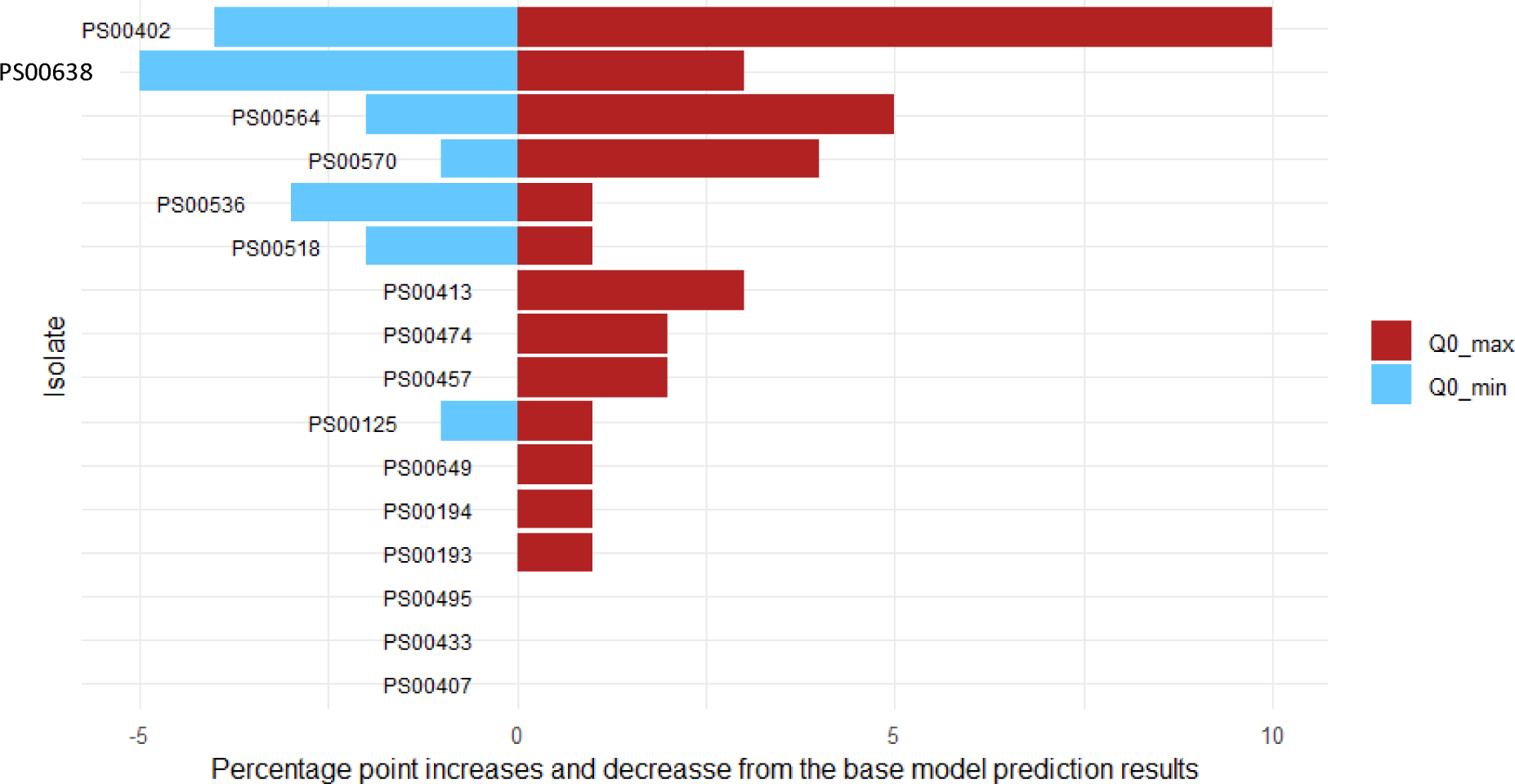
Tornado diagram for sensitivity analysis of Q0 that shows the percentage point increases (when using the maximum Q0 as the input) and decreases (when using the minimum Q0 as the input) from the base model prediction results (i.e., the percent of milk containers over 5 log_10_ CFU/ml on consumer storage day 35 when using the average Q0 as the input parameter) for 16 *B. cereus* group strains.

### Small Temperature Increases and Increased Temperature Fluctuations in Consumer Refrigerators Can Substantially Increase the Percentage of Milk Containers Predicted to Contain B. cereus Group Concentrations >10^5^ CFU/ml

What-if scenario analyses were used to evaluate the impact of four different practical scenarios on possible consumer exposure to milk with *B. cereus* group concentrations >10^5^ CFU/ml at consumer storage day 21 (representing a more realistic HTST milk shelf life) and 35 (the timing when we start to observe distinguishable variations in the percentage of milk containers containing over 10^5^ CFU/ml among *B. cereus* group strains). Our results show that a three-day delay on HTST milk product delivery from processing facility to retail without temperature abuse (scenario i) did not alter the percentage of milk containers exceeding 10^5^ CFU/ml of *B. cereus* group concentrations on consumer home storage day 21 and 35 (i.e., 2.81 and 4.13%, identical to the base model prediction). Similarly, a temperature abuse simulation of consumer transportation of HTST milk products from retail to home in a hot summer (average milk temperature of 33.5°C with an average time of 1 h; scenario ii) led to 3.06 and 4.44% milk containers exceeding 10^5^ CFU/ml on consumer home storage day 21 and 35, which are slightly higher than the base model predictions (2.81 and 4.13%). Under scenario ii), the model also predicted that three strains had higher percentages of milk containers containing *B. cereus* group concentrations >10^5^ CFU/ml compared to the baseline on consumer storage day 21. Specifically, 5 and 3% of the milk containers contaminated with isolates PS00413 and PS00433 from group IV and 5% of the milk containers contaminated with isolate PS00638 from group V had *B. cereus* group concentrations >10^5^ CFU/ml, which is higher compared to 4, 2, and 3% predicted by the base model, respectively. In contrast, two strains (PS00413 from group IV and PS00570 from group V) were predicted to exceed 10^5^ CFU/ml in a higher percentage (11% and 10%, respectively) of milk containers on consumer storage day 35, compared to the prediction by the baseline model (i.e., 10% and 6%, respectively).

The first two scenarios, which represent practical events during transportation, did not change model predictions for most, if not all, strains. However, the last two scenarios, which represent consumer behavior or ability to manage refrigeration temperature, resulted in a more profound impact. A slight temperature deviation at prolonged consumer home storage (scenario iii) had a much higher predicted impact on the percentage of milk containers containing >10^5^ CFU/ml than the two scenarios detailed above. The model predicted 4.56±3.05 (mean ± standard deviation) and 8.88±3.84 % milk containers with *B. cereus* group concentrations >10^5^ CFU/ml on consumer storage day 21 and 35, respectively (as compared to the base model prediction of 2.81 and 4.13%). Under scenario iii, 1 of 6 group IV strains and both group V strains were predicted to exceed 10^5^ CFU/ml on consumer storage day 21 in more than 10% milk containers, whereas the base model predicted that none of the strains would exceed 10^5^ CFU/ml in more than 10% milk containers. In contrast, 2 of 4 group II strains, 1 of 1 group III strain, 2 of 6 group IV strains and 2 of 2 group V strains were predicted to exceed 10^5^ CFU/ml in more than 10% milk containers on consumer storage day 35 (compared to 1 of 6 group IV strains and 1 of 2 group V strains predicted by the base model). Increasing the variability in consumer storage temperature has an even more substantial effect on the percentage of milk containers exceeding *B. cereus* group concentrations of 10^5^ CFU/ml than a consistently higher average storage temperature (scenario iii). The model predicts that 7.56±3.20 and 10.81±3.15 % milk containers had *B. cereus* group concentrations over 10^5^ CFU/ml on consumer storage day 21 and 35. Under this scenario, 1 of 4 group II strains, 1 of 6 group IV strains and both group V strains were predicted to have more than 10% of milk containers exceeding 10^5^ CFU/ml on consumer storage day 21, while 15 of 16 strains were predicted to have more than 10% of milk containers exceeding 10^5^ CFU/ml on consumer storage day 35 (compared to 0 and 2 strains predicted to have more than 10% of milk containers with over 10^5^ CFU/ml on consumer storage day 21 and 35 by the base model) (Table 5). The more profound impact from scenarios iii and iv can be explained by the fact that (i) consumer storage is the longest time in the entire supply chain, and therefore any temperature change would have a more extended effect, and (ii) changing deviation and variability increases the number of abnormal temperature conditions that exceed the growth boundaries of more strains.

**Table 5.**
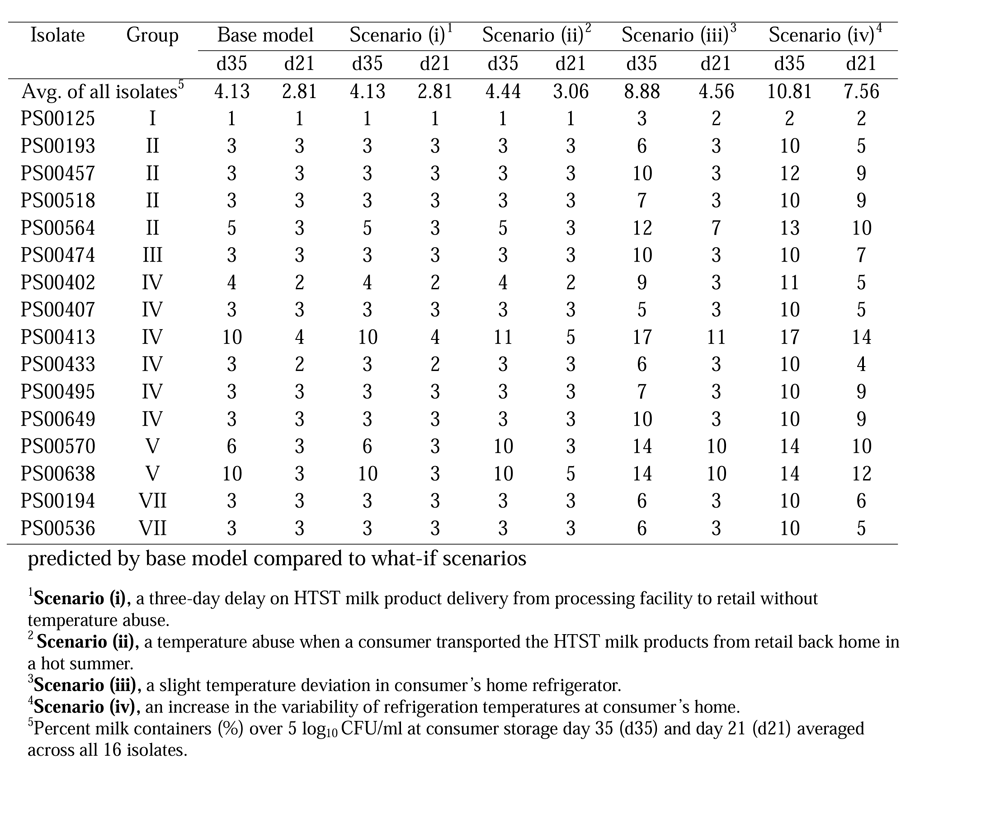
Percent milk containers over 5 log_10_ CFU/ml at consumer storage day 35 and 21.

| Isolate | Group | Base model |  | Scenario (i)<sup>1</sup> |  | Scenario (ii)<sup>2</sup> |  | Scenario (iii)<sup>3</sup> |  | Scenario (iv)<sup>4</sup> |  |
| --- | --- | --- | --- | --- | --- | --- | --- | --- | --- | --- | --- |
|  |  | d35 | d21 | d35 | d21 | d35 | d21 | d35 | d21 | d35 | d21 |
| Avg. of all isolates<sup>5</sup> |  | 4.13 | 2.81 | 4.13 | 2.81 | 4.44 | 3.06 | 8.88 | 4.56 | 10.81 | 7.56 |
| PS00125 | I | 1 | 1 | 1 | 1 | 1 | 1 | 3 | 2 | 2 | 2 |
| PS00193 | II | 3 | 3 | 3 | 3 | 3 | 3 | 6 | 3 | 10 | 5 |
| PS00457 | II | 3 | 3 | 3 | 3 | 3 | 3 | 10 | 3 | 12 | 9 |
| PS00518 | II | 3 | 3 | 3 | 3 | 3 | 3 | 7 | 3 | 10 | 9 |
| PS00564 | II | 5 | 3 | 5 | 3 | 5 | 3 | 12 | 7 | 13 | 10 |
| PS00474 | III | 3 | 3 | 3 | 3 | 3 | 3 | 10 | 3 | 10 | 7 |
| PS00402 | IV | 4 | 2 | 4 | 2 | 4 | 2 | 9 | 3 | 11 | 5 |
| PS00407 | IV | 3 | 3 | 3 | 3 | 3 | 3 | 5 | 3 | 10 | 5 |
| PS00413 | IV | 10 | 4 | 10 | 4 | 11 | 5 | 17 | 11 | 17 | 14 |
| PS00433 | IV | 3 | 2 | 3 | 2 | 3 | 3 | 6 | 3 | 10 | 4 |
| PS00495 | IV | 3 | 3 | 3 | 3 | 3 | 3 | 7 | 3 | 10 | 9 |
| PS00649 | IV | 3 | 3 | 3 | 3 | 3 | 3 | 10 | 3 | 10 | 9 |
| PS00570 | V | 6 | 3 | 6 | 3 | 10 | 3 | 14 | 10 | 14 | 10 |
| PS00638 | V | 10 | 3 | 10 | 3 | 10 | 5 | 14 | 10 | 14 | 12 |
| PS00194 | VII | 3 | 3 | 3 | 3 | 3 | 3 | 6 | 3 | 10 | 6 |
| PS00536 | VII | 3 | 3 | 3 | 3 | 3 | 3 | 6 | 3 | 10 | 5 |

predicted by base model compared to what-if scenarios
<sup>1</sup>**Scenario (i)**, a three-day delay on HTST milk product delivery from processing facility to retail without temperature abuse.
<sup>2</sup>**Scenario (ii)**, a temperature abuse when a consumer transported the HTST milk products from retail back home in a hot summer.
<sup>3</sup>**Scenario (iii)**, a slight temperature deviation in consumer's home refrigerator.
<sup>4</sup>**Scenario (iv)**, an increase in the variability of refrigeration temperatures at consumer's home.
<sup>5</sup>Percent milk containers (%) over 5 log<sub>10</sub> CFU/ml at consumer storage day 35 (d35) and day 21 (d21) averaged across all 16 isolates.

## DISCUSSION

This study collected detailed growth data for a genomically diverse set of *B. cereus* group strains and used these data to develop an exposure assessment model to facilitate industry risk assessments for diarrheal *B. cereus* group members in HTST milk. Our experimental data indicates that *B. cereus* group strains show a range of growth characteristics, including differences in minimum growth temperature, which impact the exposure through consumption of contaminated HTST milk. We specifically found that (i) *B. cereus* group strains from different phylogenetic groups differ in key cardinal values (i.e., minimum growth temperatures) and (ii) there are differences in growth parameters within a given phylogenetic group. Our data thus supports that the accuracy of *B. cereus* group exposure assessments can be improved by accounting for genetic diversity within the *B. cereus* group. Importantly, what-if scenarios run with the initial exposure assessment reported here already provide important insights into *B. cereus* group risk assessment and risk management and specifically show that absolute temperature and temperature variation during consumer storage have a large impact on predicted *B. cereus* group exposure. While our model can be useful for industry risk assessments of diarrheal *B. cereus* group members, advances in the understanding of *B. cereus* group virulence are needed to develop full risk assessment models that include public health metrics.

### Bacillus cereus Group Members Show a Range of Growth Characteristics, including Differences in Minimum Growth Temperatures and Maximum Growth Rates, which Impact the Risk Associated with the Contamination of HTST Milk

We found that the 16 tested *B. cereus* group strains showed practically relevant variations in both experimental and theoretical minimum growth temperatures. The experimentally obtained growth boundaries for most tested *B. cereuses* group strains (i.e., 14 strains) in our study were at 10°C, and two strains each had a lower or higher minimum growth temperature (8°C and 14°C). This is consistent with the finding of Lott et al. (2023) that some cytotoxic sporeformers in the *B. cereus* group may grow at abuse temperatures (e.g., 10°C), but not at lower temperatures. By comparison, one study reported that experimentally observed minimal growth temperatures of 30 diarrheal enterotoxin producing *B. cereus* group strains ranged from ≤5°C to 11°C (Dufrenne et al., 1994). Similarly, a growth study of 11 psychrotolerant *B. cereus* strains (Borge et al., 2001) reported that the lowest temperatures with observed growth (in brain heart infusion medium and commercial 1.5% UHT milk) were 4°C for one strain, 6°C for two strains and 7°C for seven strains. Guinebretière et al. (2008) reported phylogenetic group-specific minimum growth temperatures: (i) 10°C for group I (28 strains tested), (ii) 7°C for group II (33 strains tested), (iii) 15°C for group III (96 strains tested), 10°C for group IV (101 strains tested), 8°C for group V (17 strains tested), and 20°C for group VII (2 strains tested). This is generally consistent with our findings except that the group III and group VII isolates in our study had lower experimental minimum growth temperatures (i.e., 10°C) as compared to data reported by Guinebretière et al (2008).

The experimental minimum growth temperatures for all strains in our study were consistent with their theoretical minimum growth temperatures (T_min_) predicted by the secondary model. Specifically, the one group I strain with a minimum growth temperature of 14°C also had a substantially higher theoretical minimum growth temperature (T_min_) of 9.2°C compared to the other 15 strains that showed detectable growth at 10°C (T_min_ ranging from 4.26°C to 7.06°C). Notably, Carlin et al (2013) reported T_min_ for strains from group II-V are comparable to our study; however, two group VII strains in their study were reported to have substantially higher T_min_ (i.e., 12.6°C and 19.1°C) than the T_min_ in our study (i.e., 6.89°C and 7.06°C). Our growth experiment also showed that *B. cereus* group members have a wide range of estimated maximum growth rates at 22°C (0.31 ln/h to 2.02 ln/h) and 10°C (0.03 ln/h to 0.34 ln/h). This is comparable to the reported range of *B. cereus* group maximum growth rates (12 strains, growth experiment conducted in brain heart infusion supplemented with yeast extract at 2gL^−1^ and glucose at 3gL^−1^) of 0.27 h^-1^to 1.17 h^-1^ at 22°C and 0.02 h^-1^ to 0.17 h^-1^ at 10°C (Carlin et al., 2013). Overall, the high variability of growth characteristics of *B. cereus* group members, including minimum growth temperatures and maximum growth rates, suggests that it is crucial to evaluate a diverse set of strains when predicting *B. cereus* group growth. This is particularly important, as accurate predictions of the growth of different *B. cereus* strains are essential to correctly predict the exposure risk if milk is contaminated with a specific *B. cereus* group strain.

### Quantitative B. cereus Group Exposure Assessments Can Be Improved by Accounting for Genetic Diversity of B. cereus Group Members and Common HTST Milk Microbiota

Very few of the previous quantitative exposure assessments conducted in pasteurized fluid milk have addressed how the genetic diversity of *B. cereus* group members might affect human exposure (Notermans et al., 1997, Ačai et al., 2014, Lewin et al., 2019). Consistent with Carlin et al. (2013), who reported the variation of cardinal values for *B. cereus* group members with respect to their phylogenetic affiliation, our study further emphasizes the need to account for genetic variability when conducting *B. cereus* group quantitative exposure assessments. This is specifically supported by the fact that group I isolate PS00125, which has the highest T_min_ (9.22°C) in our study, was indeed predicted to have the lowest percentage of milk containers exceeding 10^5^ CFU/ml on consumer storage day 35. Importantly, model input parameters other than T_min_ (e.g., the initial physiological state of cells, Q0) may also impact exposure risk. This is supported by our exposure assessment model predictions that the risks of *B. cereus* group exposure on a given consumer storage day (especially consumer storage day 35) varies among strains and phylogenetic groups that show similar T_min_. For instance, isolates PS00413 and PS00402, both from group IV, had similar T_min_ (i.e., 4.88°C and 4.26°C); however, they showed substantial variations in another model input parameter, Q0 (i.e., 1.26E-02 and 1.91E-09). As a result, the model predicted that on consumer storage day 35, as high as 10% of the milk containers with isolate PS00413 exceeded 10^5^ CFU/ml, whereas only 4% of the milk containers with isolate PS00402 exceeded the same threshold.

Additionally, HTST milk is known to have a rich microbiota that consists of thermoduric bacteria (e.g., *Microbacterium* spp.) and gram negative bacteria associated with post-pasteurization contamination (e.g., *Pseudomonas*) (Lott, 2023). This suggests the need to account for the effect of these competing microorganisms on the growth of *B. cereus* group strains in HTST milk. Previous studies have been conducted to model the interactions between two microorganisms in the same system (Martens et al., 1999; Courtin and Rul, 2004). Similarly, future studies might be conducted to quantify the microbial interactions between *B. cereus* group strains and other microbial contaminants in the HTST milk, thus improving the accuracy of the exposure assessment model.

### What-if Scenarios Suggest That Absolute Temperature and Temperature Variation during Consumer Storage Have a Large Impact on the Predicted B. cereus Group Exposure

The exposure assessment reported here was used to run selected what-if scenarios to assess the importance of selected supply chain disruptions and changes, as well as different consumer storage temperature scenarios that may mimic practices in different locations, seasons, or infrastructure conditions. The two what-if scenarios that modelled disruption to distribution included scenarios that mimicked (i) extended storage in a distribution center (without temperature abuse) and (ii) temperature abuse during consumer transport from retail to home. Both what-if scenarios showed limited impacts on risk metrics (2.81 and 3.06% of containers that exceed 10^5^ CFU/ml, as compared to 2.81% for the base scenario on consumer storage day 21). This result is not surprising, as scenario (ii) was modelled as a relatively short time period of temperature abuse (median of 1 h; range of 14 min to 6 h). Similarly, Huang et al (2019) found that an acute temperature abuse (i.e., 35°C for 2 h) did not substantially increase the growth of foodborne pathogens (i.e., *Salmonella enterica* and *Listeria monocytogenes*) on cut fruit and radishes compared to those stored constantly at 4°C. Our results also support the recommendations for consumer time-temperature control, which is often called the “2-h rule and 1-h rule”. This rule suggests that foods should be kept outside the danger zone (4.4–60.0 °C) for less than 2 h (for temperature between 4.4 and 32.2 °C) or 1 h (for temperatures >32.2 °C) (Cho et al., 2020). Importantly, these what-if scenarios also illustrate how this model could be used to assess predicted *B. cereus* group exposures if supply chains might experience certain “worst case” disruptions (e.g., lack of refrigeration for extended time periods).

The two what-if scenarios that modelled the impact of different temperature conditions during home storage included scenarios that mimicked (i) a slight temperature deviation in a consumer’s home refrigerator, and (ii) an increase in the variability of refrigeration temperatures in a consumer’s home. These scenarios showed that even what may be considered small differences in refrigeration conditions (e.g., a change of mean temperature by 1°C) can substantially impact likely *B. cereus* group exposure. This finding aligns with our existing knowledge that prolonged temperature abuse during consumer home storage accelerates the growth of many cytotoxic *B. cereus* group isolates (Ceuppens et al., 2011). It leads to a notable occurrence, with more than 10% of milk containers exceeding 10^5^ CFU/ml on consumer home storage day 35. Of particular interest, even changes in variability of home refrigeration temperatures had substantial impacts. This type of higher variability may arise from the use of older refrigerators, more commonly prevalent in households with lower income, and/or in regions or seasons (e.g., hot summers) where rolling blackouts, i.e., planned short-term power outages, are common. Our findings are consistent with previous studies (Jofré et al., 2019) that refrigeration temperature fluctuation increases the risk of exposure to foodborne pathogens, thus negatively affecting the safety of food products. Overall, the scenarios detailed here (as well as a similar scenario that can easily be run with the code available) will help assess *B. cereus* group exposures that are likely to occur in different distribution areas that may differ in average refrigeration temperatures. Examples of distribution areas that differ in refrigeration conditions include European countries such as Germany that have a recommended refrigeration temperature of 7°C (DAAD, 2023), as compared to 4°C, the standard US refrigeration temperature recommended by FDA.

### While Our Model Will Facilitate Industry Risk Assessments for Diarrheal B. cereus Group Strains, Advances in the Understanding of B. cereus Group Virulence Are Needed to Develop Full Risk Assessments that Include Public Heath Metrics

The model reported here will provide industry with an initial tool that can facilitate risk-based food safety decision-making for products that are contaminated with low *B. cereus* group levels. For example, if an HTST fluid milk product is found to contain low levels of a specific *B. cereus* group strain, our model can be used to predict, for different time points, which proportion of products are expected to exceed a certain concentration threshold (e.g., 3 or 5 log_10_ CFU/ml). Risk managers could then use this information to inform their decision making. For example, a risk manager may decide that a product with a 21-day shelf life and a predicted 2% of containers exceeding 10^5^ CFU/ml at day 21 is considered low risk, particularly in a country with no regulatory threshold and a “low risk” supply chain (e.g., a well-developed cold chain with products typically consumed substantially before the end of the shelf life); additional considerations will likely impact these decisions (e.g., risk tolerance, lot size). In addition, this model could also be adopted to assess regulatory risk, such as the likelihood that a lot with low *B. cereus* group levels may yield a test result that exceeds a regulatory threshold, using approaches like those described previously (Chen et al., 2022). We, however, do appreciate that many users may be hesitant to use this initial model for risk management related decisions, for instance, because we currently lack the ability to predict the number of human cases that may be caused by a given lot. For example, a prediction that the risk of causing a single human case is substantially below 1% may be more valuable than the type of exposure risk that is provided as an output by our model. This type of approach has previously been used in a risk assessment that was designed to help with decision making on lots of produce that may be contaminated with *Listeria monocytogenes* (Zoellner et al., 2019). However, using the exposure assessment reported here, prediction of human illness cases that may be caused by a lot is currently impossible due to the lack of a universally applicable dose-response relationship that will allow us to predict the likelihood of *B. cereus* toxico-infection per serving and estimate the number of cases within the population (Notermans et al., 1997, Ačai et al., 2014, Lewin et al., 2019). Nevertheless, our model could be easily adapted into a full risk assessment if *B. cereus* group dose-response data become available.

## ACKNOWLEDGMENTS

This work was supported by the USDA NIFA grant 12686965, Hatch Appropriations PEN04853 and Accession 7005519, and the Multistate Capacity Grant 4666.

