## Supplement Table 1 for "Exposure assessment suggests some cytotoxic *Bacillus cereus* group genotypes can grow over 3 logs in HTST milk throughout the shelf life at temperature abuse conditions"

**Table S1.** *B. cereus* growth parameters ($\mu_{max}$, lag, $N_{max}$)[1] at 22°C, 16°C and 10°C

| Isolate | *panC* phylogenetic group | Rep[2] | Growth parameters at 10°C (or 16°C)[3] | | | Growth parameters at 22°C | | |
|---|---|---|---|---|---|---|---|---|
| | | | lag (h) | $\mu_{max}$ (ln/h) | $N_{max}$ ($log_{10}$ CFU/mL) | lag (h) | $\mu_{max}$ (ln/h) | $N_{max}$ ($log_{10}$ CFU/mL) |
| PS00125 | Group I | rep1 | 14.02 | 0.95 | 6.17 | 16.7 | 0.7 | 5.24 |
| PS00125 | Group I | rep2 | 9.12 | 0.35 | 6.28 | 14.02 | 2.02 | 5.58 |
| PS00135 | Group I | rep1 | 13.02 | 0.9 | 6.24 | 18.89 | 0.89 | 5.68 |
| PS00135 | Group I | rep2 | 12.56 | 0.9 | 6.18 | 13.7 | 0.95 | 6.03 |
| PS00193 | Group II | rep1 | 174.26 | 0.06 | 5.91 | 4.99 | 0.42 | 7.16 |
| PS00193 | Group II | rep2 | 170.88 | 0.06 | 5.88 | 10.3 | 0.83 | 6.99 |
| PS00457 | Group II | rep1 | 167.44 | 0.09 | 5.98 | 9.4 | 0.48 | 6.28 |
| PS00457 | Group II | rep2 | 101.38 | 0.07 | 6.34 | 6.71 | 0.4 | 6.35 |
| PS00518 | Group II | rep1 | 68.35 | 0.09 | 6.96 | 10.48 | 0.44 | 6.8 |
| PS00518 | Group II | rep2 | 76.84 | 0.07 | 6.59 | 18.97 | 1.41 | 6.67 |
| PS00564 | Group II | rep1 | 33.31 | 0.11 | 6.66 | 13.85 | 1.01 | 6.87 |
| PS00564 | Group II | rep2 | 62.18 | 0.14 | 6.49 | 13.28 | 0.78 | 6.75 |
| PS00474 | Group III | rep1 | 115.05 | 0.06 | 6.43 | 8.05 | 0.62 | 7.04 |
| PS00474 | Group III | rep2 | 194.05 | 0.06 | 5.01 | 3.26 | 0.4 | 7.15 |
| PS00402 | Group IV | rep1 | 180.54 | 0.34 | 6.52 | 12.77 | 0.48 | 7.09 |
| PS00402 | Group IV | rep2 | 172.2 | 0.05 | 5.01 | 7.88 | 0.46 | 6.85 |
| PS00407 | Group IV | rep1 | 41.96 | 0.04 | 6.53 | 11.96 | 0.49 | 6.86 |
| PS00407 | Group IV | rep2 | 189.76 | 0.06 | 5.41 | 8.97 | 0.48 | 6.89 |
| PS00413 | Group IV | rep1 | 27.83 | 0.11 | 7.37 | 11.24 | 0.47 | 6.99 |
| PS00413 | Group IV | rep2 | 56.56 | 0.13 | 6.51 | 4.99 | 0.39 | 7.53 |
| PS00433 | Group IV | rep1 | 81.45 | 0.05 | 6.19 | 0 | 0.31 | 7.61 |
| PS00433 | Group IV | rep2 | 122.47 | 0.06 | 6.02 | 0 | 0.33 | 7.43 |
| PS00495 | Group IV | rep1 | 72.48 | 0.03 | 6.57 | 15.32 | 0.64 | 6.99 |
| PS00495 | Group IV | rep2 | 90.45 | 0.09 | 6.36 | 7.24 | 0.58 | 7.05 |
| PS00649 | Group IV | rep1 | 100.15 | 0.08 | 6.14 | 6.85 | 0.39 | 7.14 |
| PS00649 | Group IV | rep2 | 85.6 | 0.05 | 6.46 | 11.18 | 0.62 | 6.74 |
| PS00570 | Group V | rep1 | 42.12 | 0.11 | 6.5 | 12 | 0.54 | 7.21 |
| PS00570 | Group V | rep2 | 65.76 | 0.1 | 6.72 | 5.1 | 0.38 | 7.19 |
| PS00638 | Group V | rep1 | 10.02 | 0.03 | 6.35 | 6.1 | 0.51 | 7.05 |
| PS00638 | Group V | rep2 | 106.27 | 0.09 | 5.57 | 13.86 | 0.71 | 7.21 |
| PS00194 | Group VII | rep1 | 67.98 | 0.07 | 6.71 | 22.04 | 1.08 | 6.59 |
| PS00194 | Group VII | rep2 | 110.7 | 0.11 | 6.67 | 19.47 | 0.89 | 6.74 |
| PS00536 | Group VII | rep1 | 70.25 | 0.09 | 6.82 | 21.12 | 1.78 | 6.76 |
| PS00536 | Group VII | rep2 | 86.16 | 0.06 | 6.3 | 6.32 | 0.38 | 6.39 |

[1]$\mu_{max}$ is the specific maximum growth rate in ln/h, lag is the lag time in h, $N_{max}$ is the maximum microbial population in $log_{10}$ CFU/mL.

[2]Rep is the independent biological replicate.

[3]The growth parameters ($\mu_{max}$, lag, $N_{max}$) for Isolate PS00125 and Isolate PS00135 in this column are derived from a separate 16°C growth experiment instead of the 10°C growth experiment for all the other isolates.
