## Supplement Table 2 for "Exposure assessment suggests some cytotoxic *Bacillus cereus* group genotypes can grow over 3 logs in HTST milk throughout the shelf life at temperature abuse conditions"

**Table S2.** Sensitivity analysis result of $N_{max}$. The percentage of milk containers (%) over 5 logs on consumer storage day 35 when using [1]the average, minimum, and maximum $N_{max}$ for 16 isolates are summarized.

| Isolate | *panC* phylogenetic group | Nmax_avg | Nmax_min | Nmax_max |
|---|---|---|---|---|
| PS00125 | Group I | 1 | 1 | 1 |
| PS00193 | Group II | 3 | 3 | 3 |
| PS00194 | Group VII | 3 | 3 | 3 |
| PS00402 | Group IV | 4 | 3 | 4 |
| PS00407 | Group IV | 3 | 3 | 3 |
| PS00413 | Group IV | 10 | 10 | 10 |
| PS00433 | Group IV | 3 | 3 | 3 |
| PS00457 | Group II | 3 | 3 | 3 |
| PS00474 | Group III | 3 | 3 | 3 |
| PS00495 | Group IV | 3 | 3 | 3 |
| PS00518 | Group II | 3 | 3 | 3 |
| PS00536 | Group VII | 3 | 3 | 3 |
| PS00564 | Group II | 5 | 5 | 5 |
| PS00570 | Group V | 6 | 6 | 6 |
| PS00638 | Group V | 10 | 10 | 10 |
| PS00649 | Group IV | 3 | 3 | 3 |

[1]$N_{max\_avg}$, $N_{max\_min}$ and $N_{max\_max}$ are the abbreviations for the average, minimum, and maximum $N_{max}$
