## Supplement Table 3 for "Exposure assessment suggests some cytotoxic *Bacillus cereus* group genotypes can grow over 3 logs in HTST milk throughout the shelf life at temperature abuse conditions"

**Table S3.** Growth data for 17 *B. cereus* isolates at 22, 10, 16°C.

| 22°C growth data | | | | | |
|---|---|---|---|---|---|
| Time (h) | Rep | Log Count | Isolate | *panC* phylogenetic group | Accession Number |
| 0 | rep1 | 3.44 | PS00125 | Group I | SRR23629669 |
| 1 | rep1 | 3.43 | PS00125 | Group I | SRR23629669 |
| 2 | rep1 | 3.36 | PS00125 | Group I | SRR23629669 |
| 4 | rep1 | 3.42 | PS00125 | Group I | SRR23629669 |
| 6 | rep1 | 3.46 | PS00125 | Group I | SRR23629669 |
| 8 | rep1 | 3.50 | PS00125 | Group I | SRR23629669 |
| 14 | rep1 | 3.53 | PS00125 | Group I | SRR23629669 |
| 15 | rep1 | 3.56 | PS00125 | Group I | SRR23629669 |
| 16 | rep1 | 3.57 | PS00125 | Group I | SRR23629669 |
| 18 | rep1 | 4.15 | PS00125 | Group I | SRR23629669 |
| 20 | rep1 | 4.21 | PS00125 | Group I | SRR23629669 |
| 22 | rep1 | 4.89 | PS00125 | Group I | SRR23629669 |
| 24 | rep1 | 5.23 | PS00125 | Group I | SRR23629669 |
| 27 | rep1 | 5.20 | PS00125 | Group I | SRR23629669 |
| 30 | rep1 | 5.18 | PS00125 | Group I | SRR23629669 |
| 0 | rep2 | 3.41 | PS00125 | Group I | SRR23629669 |
| 1 | rep2 | 3.41 | PS00125 | Group I | SRR23629669 |
| 2 | rep2 | 3.39 | PS00125 | Group I | SRR23629669 |
| 4 | rep2 | 3.22 | PS00125 | Group I | SRR23629669 |
| 6 | rep2 | 3.45 | PS00125 | Group I | SRR23629669 |
| 8 | rep2 | 3.59 | PS00125 | Group I | SRR23629669 |
| 14 | rep2 | 3.74 | PS00125 | Group I | SRR23629669 |
| 15 | rep2 | 4.23 | PS00125 | Group I | SRR23629669 |
| 16 | rep2 | 5.11 | PS00125 | Group I | SRR23629669 |
| 18 | rep2 | 5.21 | PS00125 | Group I | SRR23629669 |
| 20 | rep2 | 5.85 | PS00125 | Group I | SRR23629669 |
| 22 | rep2 | 6.05 | PS00125 | Group I | SRR23629669 |
| 24 | rep2 | 5.20 | PS00125 | Group I | SRR23629669 |
| 27 | rep2 | 5.27 | PS00125 | Group I | SRR23629669 |
| 30 | rep2 | 5.84 | PS00125 | Group I | SRR23629669 |
| 0 | rep1 | 3.87 | PS00135 | Group I | ASM16145v1 |
| 1 | rep1 | 3.64 | PS00135 | Group I | ASM16145v1 |
| 2 | rep1 | 3.69 | PS00135 | Group I | ASM16145v1 |
| 4 | rep1 | 3.71 | PS00135 | Group I | ASM16145v1 |
| 6 | rep1 | 3.76 | PS00135 | Group I | ASM16145v1 |
| 8 | rep1 | 3.78 | PS00135 | Group I | ASM16145v1 |
| 14 | rep1 | 3.57 | PS00135 | Group I | ASM16145v1 |
| 15 | rep1 | 3.62 | PS00135 | Group I | ASM16145v1 |
| 16 | rep1 | 3.70 | PS00135 | Group I | ASM16145v1 |
| 18 | rep1 | 4.04 | PS00135 | Group I | ASM16145v1 |
| 20 | rep1 | 4.13 | PS00135 | Group I | ASM16145v1 |

| | | | | | |
|---|---|---|---|---|---|
| 22 | rep1 | 4.99 | PS00135 | Group I | ASM16145v1 |
| 24 | rep1 | 5.20 | PS00135 | Group I | ASM16145v1 |
| 27 | rep1 | 6.16 | PS00135 | Group I | ASM16145v1 |
| 30 | rep1 | 5.26 | PS00135 | Group I | ASM16145v1 |
| 0 | rep2 | 3.60 | PS00135 | Group I | ASM16145v1 |
| 1 | rep2 | 3.61 | PS00135 | Group I | ASM16145v1 |
| 2 | rep2 | 3.54 | PS00135 | Group I | ASM16145v1 |
| 4 | rep2 | 3.70 | PS00135 | Group I | ASM16145v1 |
| 6 | rep2 | 3.76 | PS00135 | Group I | ASM16145v1 |
| 8 | rep2 | 3.75 | PS00135 | Group I | ASM16145v1 |
| 14 | rep2 | 4.01 | PS00135 | Group I | ASM16145v1 |
| 15 | rep2 | 4.14 | PS00135 | Group I | ASM16145v1 |
| 16 | rep2 | 4.91 | PS00135 | Group I | ASM16145v1 |
| 18 | rep2 | 5.15 | PS00135 | Group I | ASM16145v1 |
| 20 | rep2 | 5.91 | PS00135 | Group I | ASM16145v1 |
| 22 | rep2 | 6.11 | PS00135 | Group I | ASM16145v1 |
| 24 | rep2 | 5.93 | PS00135 | Group I | ASM16145v1 |
| 27 | rep2 | 5.98 | PS00135 | Group I | ASM16145v1 |
| 0 | rep1 | 3.58 | PS00193 | Group II | SRR5185018 |
| 1 | rep1 | 3.67 | PS00193 | Group II | SRR5185018 |
| 2 | rep1 | 3.72 | PS00193 | Group II | SRR5185018 |
| 4 | rep1 | 3.94 | PS00193 | Group II | SRR5185018 |
| 6 | rep1 | 4.06 | PS00193 | Group II | SRR5185018 |
| 8 | rep1 | 4.15 | PS00193 | Group II | SRR5185018 |
| 14 | rep1 | 5.47 | PS00193 | Group II | SRR5185018 |
| 15 | rep1 | 5.44 | PS00193 | Group II | SRR5185018 |
| 16 | rep1 | 5.65 | PS00193 | Group II | SRR5185018 |
| 18 | rep1 | 5.89 | PS00193 | Group II | SRR5185018 |
| 20 | rep1 | 6.27 | PS00193 | Group II | SRR5185018 |
| 22 | rep1 | 6.54 | PS00193 | Group II | SRR5185018 |
| 24 | rep1 | 7.07 | PS00193 | Group II | SRR5185018 |
| 27 | rep1 | 7.00 | PS00193 | Group II | SRR5185018 |
| 30 | rep1 | 7.08 | PS00193 | Group II | SRR5185018 |
| 0 | rep2 | 3.88 | PS00193 | Group II | SRR5185018 |
| 1 | rep2 | 3.95 | PS00193 | Group II | SRR5185018 |
| 2 | rep2 | 3.86 | PS00193 | Group II | SRR5185018 |
| 4 | rep2 | 3.87 | PS00193 | Group II | SRR5185018 |
| 6 | rep2 | 3.73 | PS00193 | Group II | SRR5185018 |
| 8 | rep2 | 3.84 | PS00193 | Group II | SRR5185018 |
| 14 | rep2 | 5.15 | PS00193 | Group II | SRR5185018 |
| 15 | rep2 | 5.43 | PS00193 | Group II | SRR5185018 |
| 16 | rep2 | 6.14 | PS00193 | Group II | SRR5185018 |
| 18 | rep2 | 6.32 | PS00193 | Group II | SRR5185018 |
| 20 | rep2 | 6.80 | PS00193 | Group II | SRR5185018 |
| 22 | rep2 | 6.90 | PS00193 | Group II | SRR5185018 |

| | | | | | |
|---|---|---|---|---|---|
| 24 | rep2 | 7.02 | PS00193 | Group II | SRR5185018 |
| 27 | rep2 | 6.96 | PS00193 | Group II | SRR5185018 |
| 30 | rep2 | 7.07 | PS00193 | Group II | SRR5185018 |
| 0 | rep1 | 3.79 | PS00194 | Group VII | ASM225094v2 |
| 1 | rep1 | 3.84 | PS00194 | Group VII | ASM225094v2 |
| 2 | rep1 | 3.96 | PS00194 | Group VII | ASM225094v2 |
| 4 | rep1 | 4.07 | PS00194 | Group VII | ASM225094v2 |
| 6 | rep1 | 4.17 | PS00194 | Group VII | ASM225094v2 |
| 8 | rep1 | 4.30 | PS00194 | Group VII | ASM225094v2 |
| 14 | rep1 | 3.81 | PS00194 | Group VII | ASM225094v2 |
| 15 | rep1 | 3.79 | PS00194 | Group VII | ASM225094v2 |
| 16 | rep1 | 3.80 | PS00194 | Group VII | ASM225094v2 |
| 18 | rep1 | 3.83 | PS00194 | Group VII | ASM225094v2 |
| 20 | rep1 | 3.90 | PS00194 | Group VII | ASM225094v2 |
| 22 | rep1 | 4.04 | PS00194 | Group VII | ASM225094v2 |
| 24 | rep1 | 5.07 | PS00194 | Group VII | ASM225094v2 |
| 27 | rep1 | 5.98 | PS00194 | Group VII | ASM225094v2 |
| 30 | rep1 | 6.59 | PS00194 | Group VII | ASM225094v2 |
| 0 | rep2 | 3.06 | PS00194 | Group VII | ASM225094v2 |
| 1 | rep2 | 3.12 | PS00194 | Group VII | ASM225094v2 |
| 2 | rep2 | 3.63 | PS00194 | Group VII | ASM225094v2 |
| 4 | rep2 | 3.44 | PS00194 | Group VII | ASM225094v2 |
| 6 | rep2 | 3.53 | PS00194 | Group VII | ASM225094v2 |
| 8 | rep2 | 3.76 | PS00194 | Group VII | ASM225094v2 |
| 14 | rep2 | 3.66 | PS00194 | Group VII | ASM225094v2 |
| 15 | rep2 | 3.65 | PS00194 | Group VII | ASM225094v2 |
| 16 | rep2 | 3.59 | PS00194 | Group VII | ASM225094v2 |
| 18 | rep2 | 3.70 | PS00194 | Group VII | ASM225094v2 |
| 20 | rep2 | 4.19 | PS00194 | Group VII | ASM225094v2 |
| 22 | rep2 | 4.29 | PS00194 | Group VII | ASM225094v2 |
| 24 | rep2 | 5.14 | PS00194 | Group VII | ASM225094v2 |
| 27 | rep2 | 6.48 | PS00194 | Group VII | ASM225094v2 |
| 30 | rep2 | 6.61 | PS00194 | Group VII | ASM225094v2 |
| 0 | rep1 | 3.60 | PS00402 | Group IV | SRR23629691 |
| 1 | rep1 | 3.69 | PS00402 | Group IV | SRR23629691 |
| 2 | rep1 | 3.92 | PS00402 | Group IV | SRR23629691 |
| 4 | rep1 | 4.11 | PS00402 | Group IV | SRR23629691 |
| 6 | rep1 | 4.25 | PS00402 | Group IV | SRR23629691 |
| 8 | rep1 | 4.32 | PS00402 | Group IV | SRR23629691 |
| 14 | rep1 | 4.51 | PS00402 | Group IV | SRR23629691 |
| 15 | rep1 | 4.51 | PS00402 | Group IV | SRR23629691 |
| 16 | rep1 | 4.67 | PS00402 | Group IV | SRR23629691 |
| 18 | rep1 | 5.13 | PS00402 | Group IV | SRR23629691 |
| 20 | rep1 | 5.27 | PS00402 | Group IV | SRR23629691 |
| 22 | rep1 | 5.95 | PS00402 | Group IV | SRR23629691 |

| | | | | | |
|---|---|---|---|---|---|
| 24 | rep1 | 6.26 | PS00402 | Group IV | SRR23629691 |
| 27 | rep1 | 6.89 | PS00402 | Group IV | SRR23629691 |
| 30 | rep1 | 6.82 | PS00402 | Group IV | SRR23629691 |
| 32 | rep1 | 7.07 | PS00402 | Group IV | SRR23629691 |
| 0 | rep2 | 3.77 | PS00402 | Group IV | SRR23629691 |
| 1 | rep2 | 3.78 | PS00402 | Group IV | SRR23629691 |
| 2 | rep2 | 3.80 | PS00402 | Group IV | SRR23629691 |
| 4 | rep2 | 3.82 | PS00402 | Group IV | SRR23629691 |
| 6 | rep2 | 3.83 | PS00402 | Group IV | SRR23629691 |
| 8 | rep2 | 4.00 | PS00402 | Group IV | SRR23629691 |
| 14 | rep2 | 5.00 | PS00402 | Group IV | SRR23629691 |
| 15 | rep2 | 5.11 | PS00402 | Group IV | SRR23629691 |
| 16 | rep2 | 5.42 | PS00402 | Group IV | SRR23629691 |
| 18 | rep2 | 5.83 | PS00402 | Group IV | SRR23629691 |
| 20 | rep2 | 6.23 | PS00402 | Group IV | SRR23629691 |
| 22 | rep2 | 6.27 | PS00402 | Group IV | SRR23629691 |
| 24 | rep2 | 6.41 | PS00402 | Group IV | SRR23629691 |
| 27 | rep2 | 6.77 | PS00402 | Group IV | SRR23629691 |
| 30 | rep2 | 6.87 | PS00402 | Group IV | SRR23629691 |
| 32 | rep2 | 6.93 | PS00402 | Group IV | SRR23629691 |
| 0 | rep1 | 3.74 | PS00407 | Group IV | SRR13988985 |
| 1 | rep1 | 3.79 | PS00407 | Group IV | SRR13988985 |
| 2 | rep1 | 3.87 | PS00407 | Group IV | SRR13988985 |
| 4 | rep1 | 3.97 | PS00407 | Group IV | SRR13988985 |
| 6 | rep1 | 4.08 | PS00407 | Group IV | SRR13988985 |
| 8 | rep1 | 4.20 | PS00407 | Group IV | SRR13988985 |
| 14 | rep1 | 4.62 | PS00407 | Group IV | SRR13988985 |
| 15 | rep1 | 4.71 | PS00407 | Group IV | SRR13988985 |
| 16 | rep1 | 4.75 | PS00407 | Group IV | SRR13988985 |
| 18 | rep1 | 5.05 | PS00407 | Group IV | SRR13988985 |
| 20 | rep1 | 5.59 | PS00407 | Group IV | SRR13988985 |
| 22 | rep1 | 5.82 | PS00407 | Group IV | SRR13988985 |
| 24 | rep1 | 6.68 | PS00407 | Group IV | SRR13988985 |
| 27 | rep1 | 6.66 | PS00407 | Group IV | SRR13988985 |
| 30 | rep1 | 6.73 | PS00407 | Group IV | SRR13988985 |
| 0 | rep2 | 3.67 | PS00407 | Group IV | SRR13988985 |
| 1 | rep2 | 3.66 | PS00407 | Group IV | SRR13988985 |
| 2 | rep2 | 3.66 | PS00407 | Group IV | SRR13988985 |
| 4 | rep2 | 3.72 | PS00407 | Group IV | SRR13988985 |
| 6 | rep2 | 3.80 | PS00407 | Group IV | SRR13988985 |
| 8 | rep2 | 3.86 | PS00407 | Group IV | SRR13988985 |
| 14 | rep2 | 4.75 | PS00407 | Group IV | SRR13988985 |
| 15 | rep2 | 4.99 | PS00407 | Group IV | SRR13988985 |
| 16 | rep2 | 5.14 | PS00407 | Group IV | SRR13988985 |
| 18 | rep2 | 5.56 | PS00407 | Group IV | SRR13988985 |

| | | | | | |
|---|---|---|---|---|---|
| 20 | rep2 | 5.93 | PS00407 | Group IV | SRR13988985 |
| 22 | rep2 | 6.14 | PS00407 | Group IV | SRR13988985 |
| 24 | rep2 | 6.68 | PS00407 | Group IV | SRR13988985 |
| 27 | rep2 | 6.86 | PS00407 | Group IV | SRR13988985 |
| 30 | rep2 | 6.78 | PS00407 | Group IV | SRR13988985 |
| 0 | rep1 | 3.91 | PS00413 | Group IV | SRR13988908 |
| 1 | rep1 | 3.94 | PS00413 | Group IV | SRR13988908 |
| 2 | rep1 | 4.01 | PS00413 | Group IV | SRR13988908 |
| 4 | rep1 | 4.11 | PS00413 | Group IV | SRR13988908 |
| 6 | rep1 | 4.20 | PS00413 | Group IV | SRR13988908 |
| 8 | rep1 | 4.32 | PS00413 | Group IV | SRR13988908 |
| 14 | rep1 | 4.76 | PS00413 | Group IV | SRR13988908 |
| 15 | rep1 | 4.75 | PS00413 | Group IV | SRR13988908 |
| 16 | rep1 | 4.99 | PS00413 | Group IV | SRR13988908 |
| 18 | rep1 | 5.60 | PS00413 | Group IV | SRR13988908 |
| 20 | rep1 | 5.63 | PS00413 | Group IV | SRR13988908 |
| 22 | rep1 | 6.54 | PS00413 | Group IV | SRR13988908 |
| 24 | rep1 | 6.27 | PS00413 | Group IV | SRR13988908 |
| 27 | rep1 | 6.91 | PS00413 | Group IV | SRR13988908 |
| 30 | rep1 | 6.91 | PS00413 | Group IV | SRR13988908 |
| 0 | rep2 | 2.98 | PS00413 | Group IV | SRR13988908 |
| 1 | rep2 | 3.26 | PS00413 | Group IV | SRR13988908 |
| 2 | rep2 | 3.69 | PS00413 | Group IV | SRR13988908 |
| 4 | rep2 | 3.91 | PS00413 | Group IV | SRR13988908 |
| 6 | rep2 | 3.84 | PS00413 | Group IV | SRR13988908 |
| 8 | rep2 | 4.02 | PS00413 | Group IV | SRR13988908 |
| 14 | rep2 | 4.69 | PS00413 | Group IV | SRR13988908 |
| 15 | rep2 | 4.76 | PS00413 | Group IV | SRR13988908 |
| 16 | rep2 | 5.17 | PS00413 | Group IV | SRR13988908 |
| 18 | rep2 | 5.67 | PS00413 | Group IV | SRR13988908 |
| 20 | rep2 | 5.92 | PS00413 | Group IV | SRR13988908 |
| 22 | rep2 | 5.98 | PS00413 | Group IV | SRR13988908 |
| 24 | rep2 | 6.91 | PS00413 | Group IV | SRR13988908 |
| 27 | rep2 | 7.06 | PS00413 | Group IV | SRR13988908 |
| 30 | rep2 | 7.15 | PS00413 | Group IV | SRR13988908 |
| 0 | rep1 | 3.76 | PS00433 | Group IV | SRR13988883 |
| 1 | rep1 | 3.78 | PS00433 | Group IV | SRR13988883 |
| 2 | rep1 | 3.85 | PS00433 | Group IV | SRR13988883 |
| 4 | rep1 | 4.08 | PS00433 | Group IV | SRR13988883 |
| 6 | rep1 | 4.22 | PS00433 | Group IV | SRR13988883 |
| 8 | rep1 | 4.33 | PS00433 | Group IV | SRR13988883 |
| 14 | rep1 | 5.32 | PS00433 | Group IV | SRR13988883 |
| 15 | rep1 | 5.61 | PS00433 | Group IV | SRR13988883 |
| 16 | rep1 | 5.45 | PS00433 | Group IV | SRR13988883 |
| 18 | rep1 | 5.92 | PS00433 | Group IV | SRR13988883 |

| | | | | | |
|---|---|---|---|---|---|
| 20 | rep1 | 6.13 | PS00433 | Group IV | SRR13988883 |
| 22 | rep1 | 6.69 | PS00433 | Group IV | SRR13988883 |
| 24 | rep1 | 6.96 | PS00433 | Group IV | SRR13988883 |
| 27 | rep1 | 7.13 | PS00433 | Group IV | SRR13988883 |
| 30 | rep1 | 7.17 | PS00433 | Group IV | SRR13988883 |
| 0 | rep2 | 3.31 | PS00433 | Group IV | SRR13988883 |
| 1 | rep2 | 3.66 | PS00433 | Group IV | SRR13988883 |
| 2 | rep2 | 3.78 | PS00433 | Group IV | SRR13988883 |
| 4 | rep2 | 3.82 | PS00433 | Group IV | SRR13988883 |
| 6 | rep2 | 4.01 | PS00433 | Group IV | SRR13988883 |
| 8 | rep2 | 5.19 | PS00433 | Group IV | SRR13988883 |
| 14 | rep2 | 5.32 | PS00433 | Group IV | SRR13988883 |
| 15 | rep2 | 5.33 | PS00433 | Group IV | SRR13988883 |
| 16 | rep2 | 5.54 | PS00433 | Group IV | SRR13988883 |
| 18 | rep2 | 5.88 | PS00433 | Group IV | SRR13988883 |
| 20 | rep2 | 6.04 | PS00433 | Group IV | SRR13988883 |
| 22 | rep2 | 6.53 | PS00433 | Group IV | SRR13988883 |
| 24 | rep2 | 7.16 | PS00433 | Group IV | SRR13988883 |
| 27 | rep2 | 7.03 | PS00433 | Group IV | SRR13988883 |
| 30 | rep2 | 7.12 | PS00433 | Group IV | SRR13988883 |
| 0 | rep1 | 3.12 | PS00457 | Group II | SRR13988862 |
| 1 | rep1 | 3.19 | PS00457 | Group II | SRR13988862 |
| 2 | rep1 | 3.35 | PS00457 | Group II | SRR13988862 |
| 4 | rep1 | 3.57 | PS00457 | Group II | SRR13988862 |
| 6 | rep1 | 3.65 | PS00457 | Group II | SRR13988862 |
| 8 | rep1 | 3.72 | PS00457 | Group II | SRR13988862 |
| 14 | rep1 | 4.45 | PS00457 | Group II | SRR13988862 |
| 15 | rep1 | 4.58 | PS00457 | Group II | SRR13988862 |
| 16 | rep1 | 4.68 | PS00457 | Group II | SRR13988862 |
| 18 | rep1 | 4.86 | PS00457 | Group II | SRR13988862 |
| 20 | rep1 | 5.26 | PS00457 | Group II | SRR13988862 |
| 22 | rep1 | 6.26 | PS00457 | Group II | SRR13988862 |
| 24 | rep1 | 6.34 | PS00457 | Group II | SRR13988862 |
| 27 | rep1 | 6.17 | PS00457 | Group II | SRR13988862 |
| 30 | rep1 | 6.06 | PS00457 | Group II | SRR13988862 |
| 0 | rep2 | 3.02 | PS00457 | Group II | SRR13988862 |
| 1 | rep2 | 3.15 | PS00457 | Group II | SRR13988862 |
| 2 | rep2 | 3.33 | PS00457 | Group II | SRR13988862 |
| 4 | rep2 | 3.48 | PS00457 | Group II | SRR13988862 |
| 6 | rep2 | 3.67 | PS00457 | Group II | SRR13988862 |
| 8 | rep2 | 3.78 | PS00457 | Group II | SRR13988862 |
| 14 | rep2 | 4.53 | PS00457 | Group II | SRR13988862 |
| 15 | rep2 | 4.61 | PS00457 | Group II | SRR13988862 |
| 16 | rep2 | 4.71 | PS00457 | Group II | SRR13988862 |
| 18 | rep2 | 4.90 | PS00457 | Group II | SRR13988862 |

| | | | | | |
|---|---|---|---|---|---|
| 20 | rep2 | 5.23 | PS00457 | Group II | SRR13988862 |
| 22 | rep2 | 6.21 | PS00457 | Group II | SRR13988862 |
| 24 | rep2 | 6.32 | PS00457 | Group II | SRR13988862 |
| 27 | rep2 | 6.16 | PS00457 | Group II | SRR13988862 |
| 30 | rep2 | 6.12 | PS00457 | Group II | SRR13988862 |
| 0 | rep1 | 3.61 | PS00457 | Group II | SRR13988862 |
| 1 | rep1 | 3.65 | PS00457 | Group II | SRR13988862 |
| 2 | rep1 | 3.82 | PS00457 | Group II | SRR13988862 |
| 4 | rep1 | 4.05 | PS00457 | Group II | SRR13988862 |
| 6 | rep1 | 4.16 | PS00457 | Group II | SRR13988862 |
| 8 | rep1 | 4.30 | PS00457 | Group II | SRR13988862 |
| 14 | rep1 | 4.99 | PS00457 | Group II | SRR13988862 |
| 15 | rep1 | 5.65 | PS00457 | Group II | SRR13988862 |
| 16 | rep1 | 6.04 | PS00457 | Group II | SRR13988862 |
| 18 | rep1 | 6.67 | PS00457 | Group II | SRR13988862 |
| 20 | rep1 | 6.77 | PS00457 | Group II | SRR13988862 |
| 22 | rep1 | 6.86 | PS00457 | Group II | SRR13988862 |
| 24 | rep1 | 7.04 | PS00457 | Group II | SRR13988862 |
| 27 | rep1 | 7.01 | PS00457 | Group II | SRR13988862 |
| 30 | rep1 | 7.02 | PS00457 | Group II | SRR13988862 |
| 0 | rep2 | 3.96 | PS00457 | Group II | SRR13988862 |
| 1 | rep2 | 4.13 | PS00457 | Group II | SRR13988862 |
| 2 | rep2 | 4.34 | PS00457 | Group II | SRR13988862 |
| 4 | rep2 | 4.53 | PS00457 | Group II | SRR13988862 |
| 6 | rep2 | 4.64 | PS00457 | Group II | SRR13988862 |
| 8 | rep2 | 4.83 | PS00457 | Group II | SRR13988862 |
| 14 | rep2 | 5.88 | PS00457 | Group II | SRR13988862 |
| 15 | rep2 | 6.00 | PS00457 | Group II | SRR13988862 |
| 16 | rep2 | 6.20 | PS00457 | Group II | SRR13988862 |
| 18 | rep2 | 6.63 | PS00457 | Group II | SRR13988862 |
| 20 | rep2 | 6.87 | PS00457 | Group II | SRR13988862 |
| 22 | rep2 | 6.85 | PS00457 | Group II | SRR13988862 |
| 24 | rep2 | 6.94 | PS00457 | Group II | SRR13988862 |
| 27 | rep2 | 7.17 | PS00457 | Group II | SRR13988862 |
| 0 | rep1 | 3.68 | PS00495 | Group IV | SRR13988829 |
| 1 | rep1 | 3.71 | PS00495 | Group IV | SRR13988829 |
| 2 | rep1 | 3.80 | PS00495 | Group IV | SRR13988829 |
| 4 | rep1 | 4.00 | PS00495 | Group IV | SRR13988829 |
| 6 | rep1 | 4.16 | PS00495 | Group IV | SRR13988829 |
| 8 | rep1 | 4.36 | PS00495 | Group IV | SRR13988829 |
| 14 | rep1 | 4.18 | PS00495 | Group IV | SRR13988829 |
| 15 | rep1 | 4.26 | PS00495 | Group IV | SRR13988829 |
| 16 | rep1 | 4.30 | PS00495 | Group IV | SRR13988829 |
| 18 | rep1 | 4.94 | PS00495 | Group IV | SRR13988829 |
| 20 | rep1 | 5.10 | PS00495 | Group IV | SRR13988829 |

| | | | | | |
|---|---|---|---|---|---|
| 22 | rep1 | 5.69 | PS00495 | Group IV | SRR13988829 |
| 24 | rep1 | 6.38 | PS00495 | Group IV | SRR13988829 |
| 27 | rep1 | 6.92 | PS00495 | Group IV | SRR13988829 |
| 30 | rep1 | 6.85 | PS00495 | Group IV | SRR13988829 |
| 0 | rep2 | 3.71 | PS00495 | Group IV | SRR13988829 |
| 1 | rep2 | 3.78 | PS00495 | Group IV | SRR13988829 |
| 2 | rep2 | 3.80 | PS00495 | Group IV | SRR13988829 |
| 4 | rep2 | 3.92 | PS00495 | Group IV | SRR13988829 |
| 6 | rep2 | 3.93 | PS00495 | Group IV | SRR13988829 |
| 8 | rep2 | 4.04 | PS00495 | Group IV | SRR13988829 |
| 14 | rep2 | 5.68 | PS00495 | Group IV | SRR13988829 |
| 15 | rep2 | 5.72 | PS00495 | Group IV | SRR13988829 |
| 16 | rep2 | 5.91 | PS00495 | Group IV | SRR13988829 |
| 18 | rep2 | 6.20 | PS00495 | Group IV | SRR13988829 |
| 20 | rep2 | 6.75 | PS00495 | Group IV | SRR13988829 |
| 22 | rep2 | 6.97 | PS00495 | Group IV | SRR13988829 |
| 24 | rep2 | 6.98 | PS00495 | Group IV | SRR13988829 |
| 27 | rep2 | 7.04 | PS00495 | Group IV | SRR13988829 |
| 0 | rep1 | 3.53 | PS00518 | Group II | SRR13989003 |
| 1 | rep1 | 3.96 | PS00518 | Group II | SRR13989003 |
| 2 | rep1 | 4.14 | PS00518 | Group II | SRR13989003 |
| 4 | rep1 | 4.20 | PS00518 | Group II | SRR13989003 |
| 6 | rep1 | 4.23 | PS00518 | Group II | SRR13989003 |
| 8 | rep1 | 4.28 | PS00518 | Group II | SRR13989003 |
| 14 | rep1 | 4.71 | PS00518 | Group II | SRR13989003 |
| 15 | rep1 | 4.82 | PS00518 | Group II | SRR13989003 |
| 16 | rep1 | 4.81 | PS00518 | Group II | SRR13989003 |
| 18 | rep1 | 5.74 | PS00518 | Group II | SRR13989003 |
| 20 | rep1 | 5.96 | PS00518 | Group II | SRR13989003 |
| 22 | rep1 | 6.18 | PS00518 | Group II | SRR13989003 |
| 24 | rep1 | 6.12 | PS00518 | Group II | SRR13989003 |
| 27 | rep1 | 6.73 | PS00518 | Group II | SRR13989003 |
| 30 | rep1 | 6.76 | PS00518 | Group II | SRR13989003 |
| 0 | rep2 | 3.29 | PS00518 | Group II | SRR13989003 |
| 1 | rep2 | 3.12 | PS00518 | Group II | SRR13989003 |
| 2 | rep2 | 3.14 | PS00518 | Group II | SRR13989003 |
| 4 | rep2 | 3.24 | PS00518 | Group II | SRR13989003 |
| 6 | rep2 | 3.14 | PS00518 | Group II | SRR13989003 |
| 8 | rep2 | 3.31 | PS00518 | Group II | SRR13989003 |
| 14 | rep2 | 2.89 | PS00518 | Group II | SRR13989003 |
| 15 | rep2 | 3.08 | PS00518 | Group II | SRR13989003 |
| 16 | rep2 | 3.20 | PS00518 | Group II | SRR13989003 |
| 18 | rep2 | 3.43 | PS00518 | Group II | SRR13989003 |
| 20 | rep2 | 3.99 | PS00518 | Group II | SRR13989003 |
| 22 | rep2 | 4.78 | PS00518 | Group II | SRR13989003 |

| | | | | | |
|---|---|---|---|---|---|
| 24 | rep2 | 6.30 | PS00518 | Group II | SRR13989003 |
| 27 | rep2 | 6.70 | PS00518 | Group II | SRR13989003 |
| 30 | rep2 | 6.58 | PS00518 | Group II | SRR13989003 |
| 0 | rep1 | 3.56 | PS00536 | Group VII | SRR13988990 |
| 1 | rep1 | 3.54 | PS00536 | Group VII | SRR13988990 |
| 2 | rep1 | 3.59 | PS00536 | Group VII | SRR13988990 |
| 4 | rep1 | 3.79 | PS00536 | Group VII | SRR13988990 |
| 6 | rep1 | 3.75 | PS00536 | Group VII | SRR13988990 |
| 8 | rep1 | 3.88 | PS00536 | Group VII | SRR13988990 |
| 14 | rep1 | 3.73 | PS00536 | Group VII | SRR13988990 |
| 15 | rep1 | 3.77 | PS00536 | Group VII | SRR13988990 |
| 16 | rep1 | 3.78 | PS00536 | Group VII | SRR13988990 |
| 18 | rep1 | 3.78 | PS00536 | Group VII | SRR13988990 |
| 20 | rep1 | 3.87 | PS00536 | Group VII | SRR13988990 |
| 22 | rep1 | 4.43 | PS00536 | Group VII | SRR13988990 |
| 24 | rep1 | 5.94 | PS00536 | Group VII | SRR13988990 |
| 27 | rep1 | 6.32 | PS00536 | Group VII | SRR13988990 |
| 30 | rep1 | 6.92 | PS00536 | Group VII | SRR13988990 |
| 32 | rep1 | 7.01 | PS00536 | Group VII | SRR13988990 |
| 0 | rep2 | 3.01 | PS00536 | Group VII | SRR13988990 |
| 1 | rep2 | 3.05 | PS00536 | Group VII | SRR13988990 |
| 2 | rep2 | 3.09 | PS00536 | Group VII | SRR13988990 |
| 4 | rep2 | 3.25 | PS00536 | Group VII | SRR13988990 |
| 6 | rep2 | 3.52 | PS00536 | Group VII | SRR13988990 |
| 8 | rep2 | 3.76 | PS00536 | Group VII | SRR13988990 |
| 14 | rep2 | 4.36 | PS00536 | Group VII | SRR13988990 |
| 15 | rep2 | 4.52 | PS00536 | Group VII | SRR13988990 |
| 16 | rep2 | 4.58 | PS00536 | Group VII | SRR13988990 |
| 18 | rep2 | 4.65 | PS00536 | Group VII | SRR13988990 |
| 20 | rep2 | 5.23 | PS00536 | Group VII | SRR13988990 |
| 22 | rep2 | 5.94 | PS00536 | Group VII | SRR13988990 |
| 24 | rep2 | 6.15 | PS00536 | Group VII | SRR13988990 |
| 27 | rep2 | 6.20 | PS00536 | Group VII | SRR13988990 |
| 30 | rep2 | 6.14 | PS00536 | Group VII | SRR13988990 |
| 0 | rep1 | 3.94 | PS00564 | Group II | SRR13988960 |
| 1 | rep1 | 4.08 | PS00564 | Group II | SRR13988960 |
| 2 | rep1 | 4.01 | PS00564 | Group II | SRR13988960 |
| 4 | rep1 | 3.84 | PS00564 | Group II | SRR13988960 |
| 6 | rep1 | 3.81 | PS00564 | Group II | SRR13988960 |
| 8 | rep1 | 3.88 | PS00564 | Group II | SRR13988960 |
| 14 | rep1 | 4.06 | PS00564 | Group II | SRR13988960 |
| 15 | rep1 | 4.62 | PS00564 | Group II | SRR13988960 |
| 16 | rep1 | 4.94 | PS00564 | Group II | SRR13988960 |
| 18 | rep1 | 5.81 | PS00564 | Group II | SRR13988960 |
| 20 | rep1 | 6.34 | PS00564 | Group II | SRR13988960 |

| | | | | | |
|---|---|---|---|---|---|
| 22 | rep1 | 6.68 | PS00564 | Group II | SRR13988960 |
| 24 | rep1 | 6.98 | PS00564 | Group II | SRR13988960 |
| 27 | rep1 | 6.89 | PS00564 | Group II | SRR13988960 |
| 30 | rep1 | 6.86 | PS00564 | Group II | SRR13988960 |
| 0 | rep2 | 3.51 | PS00564 | Group II | SRR13988960 |
| 1 | rep2 | 3.53 | PS00564 | Group II | SRR13988960 |
| 2 | rep2 | 3.56 | PS00564 | Group II | SRR13988960 |
| 4 | rep2 | 3.57 | PS00564 | Group II | SRR13988960 |
| 6 | rep2 | 3.55 | PS00564 | Group II | SRR13988960 |
| 8 | rep2 | 3.58 | PS00564 | Group II | SRR13988960 |
| 14 | rep2 | 3.87 | PS00564 | Group II | SRR13988960 |
| 15 | rep2 | 4.25 | PS00564 | Group II | SRR13988960 |
| 16 | rep2 | 4.61 | PS00564 | Group II | SRR13988960 |
| 18 | rep2 | 5.13 | PS00564 | Group II | SRR13988960 |
| 20 | rep2 | 5.74 | PS00564 | Group II | SRR13988960 |
| 22 | rep2 | 6.24 | PS00564 | Group II | SRR13988960 |
| 24 | rep2 | 6.72 | PS00564 | Group II | SRR13988960 |
| 27 | rep2 | 6.60 | PS00564 | Group II | SRR13988960 |
| 30 | rep2 | 6.83 | PS00564 | Group II | SRR13988960 |
| 0 | rep1 | 3.53 | PS00570 | Group V | SRR13988955 |
| 1 | rep1 | 3.58 | PS00570 | Group V | SRR13988955 |
| 2 | rep1 | 3.79 | PS00570 | Group V | SRR13988955 |
| 4 | rep1 | 4.01 | PS00570 | Group V | SRR13988955 |
| 6 | rep1 | 4.22 | PS00570 | Group V | SRR13988955 |
| 8 | rep1 | 4.30 | PS00570 | Group V | SRR13988955 |
| 14 | rep1 | 4.58 | PS00570 | Group V | SRR13988955 |
| 15 | rep1 | 4.53 | PS00570 | Group V | SRR13988955 |
| 16 | rep1 | 4.98 | PS00570 | Group V | SRR13988955 |
| 18 | rep1 | 5.27 | PS00570 | Group V | SRR13988955 |
| 20 | rep1 | 5.57 | PS00570 | Group V | SRR13988955 |
| 22 | rep1 | 6.03 | PS00570 | Group V | SRR13988955 |
| 24 | rep1 | 6.99 | PS00570 | Group V | SRR13988955 |
| 27 | rep1 | 6.98 | PS00570 | Group V | SRR13988955 |
| 30 | rep1 | 7.10 | PS00570 | Group V | SRR13988955 |
| 0 | rep2 | 3.98 | PS00570 | Group V | SRR13988955 |
| 1 | rep2 | 4.11 | PS00570 | Group V | SRR13988955 |
| 2 | rep2 | 4.21 | PS00570 | Group V | SRR13988955 |
| 4 | rep2 | 4.33 | PS00570 | Group V | SRR13988955 |
| 6 | rep2 | 4.45 | PS00570 | Group V | SRR13988955 |
| 8 | rep2 | 4.52 | PS00570 | Group V | SRR13988955 |
| 14 | rep2 | 5.68 | PS00570 | Group V | SRR13988955 |
| 15 | rep2 | 5.68 | PS00570 | Group V | SRR13988955 |
| 16 | rep2 | 5.79 | PS00570 | Group V | SRR13988955 |
| 18 | rep2 | 6.21 | PS00570 | Group V | SRR13988955 |
| 20 | rep2 | 6.47 | PS00570 | Group V | SRR13988955 |

| | | | | | |
|---|---|---|---|---|---|
| 22 | rep2 | 6.64 | PS00570 | Group V | SRR13988955 |
| 24 | rep2 | 6.86 | PS00570 | Group V | SRR13988955 |
| 27 | rep2 | 7.14 | PS00570 | Group V | SRR13988955 |
| 30 | rep2 | 7.12 | PS00570 | Group V | SRR13988955 |
| 0 | rep1 | 3.78 | PS00638 | Group V | SRR13988845 |
| 1 | rep1 | 3.80 | PS00638 | Group V | SRR13988845 |
| 2 | rep1 | 3.86 | PS00638 | Group V | SRR13988845 |
| 4 | rep1 | 3.96 | PS00638 | Group V | SRR13988845 |
| 6 | rep1 | 4.04 | PS00638 | Group V | SRR13988845 |
| 8 | rep1 | 4.19 | PS00638 | Group V | SRR13988845 |
| 14 | rep1 | 5.68 | PS00638 | Group V | SRR13988845 |
| 15 | rep1 | 5.75 | PS00638 | Group V | SRR13988845 |
| 16 | rep1 | 6.09 | PS00638 | Group V | SRR13988845 |
| 18 | rep1 | 6.34 | PS00638 | Group V | SRR13988845 |
| 20 | rep1 | 6.40 | PS00638 | Group V | SRR13988845 |
| 22 | rep1 | 6.84 | PS00638 | Group V | SRR13988845 |
| 24 | rep1 | 6.99 | PS00638 | Group V | SRR13988845 |
| 27 | rep1 | 7.10 | PS00638 | Group V | SRR13988845 |
| 30 | rep1 | 7.07 | PS00638 | Group V | SRR13988845 |
| 0 | rep2 | 4.00 | PS00638 | Group V | SRR13988845 |
| 1 | rep2 | 4.28 | PS00638 | Group V | SRR13988845 |
| 2 | rep2 | 4.31 | PS00638 | Group V | SRR13988845 |
| 4 | rep2 | 4.19 | PS00638 | Group V | SRR13988845 |
| 6 | rep2 | 4.62 | PS00638 | Group V | SRR13988845 |
| 8 | rep2 | 4.83 | PS00638 | Group V | SRR13988845 |
| 14 | rep2 | 4.58 | PS00638 | Group V | SRR13988845 |
| 15 | rep2 | 4.92 | PS00638 | Group V | SRR13988845 |
| 16 | rep2 | 5.12 | PS00638 | Group V | SRR13988845 |
| 18 | rep2 | 5.67 | PS00638 | Group V | SRR13988845 |
| 20 | rep2 | 6.19 | PS00638 | Group V | SRR13988845 |
| 22 | rep2 | 6.68 | PS00638 | Group V | SRR13988845 |
| 24 | rep2 | 7.08 | PS00638 | Group V | SRR13988845 |
| 27 | rep2 | 7.16 | PS00638 | Group V | SRR13988845 |
| 0 | rep1 | 3.56 | PS00649 | Group IV | SRR23629673 |
| 1 | rep1 | 3.63 | PS00649 | Group IV | SRR23629673 |
| 2 | rep1 | 3.78 | PS00649 | Group IV | SRR23629673 |
| 4 | rep1 | 3.94 | PS00649 | Group IV | SRR23629673 |
| 6 | rep1 | 4.04 | PS00649 | Group IV | SRR23629673 |
| 8 | rep1 | 4.21 | PS00649 | Group IV | SRR23629673 |
| 14 | rep1 | 4.76 | PS00649 | Group IV | SRR23629673 |
| 15 | rep1 | 4.92 | PS00649 | Group IV | SRR23629673 |
| 16 | rep1 | 5.11 | PS00649 | Group IV | SRR23629673 |
| 18 | rep1 | 5.63 | PS00649 | Group IV | SRR23629673 |
| 20 | rep1 | 6.17 | PS00649 | Group IV | SRR23629673 |
| 22 | rep1 | 6.39 | PS00649 | Group IV | SRR23629673 |

| | | | | | |
|---|---|---|---|---|---|
| 24 | rep1 | 6.30 | PS00649 | Group IV | SRR23629673 |
| 27 | rep1 | 6.71 | PS00649 | Group IV | SRR23629673 |
| 30 | rep1 | 7.08 | PS00649 | Group IV | SRR23629673 |
| 0 | rep2 | 3.11 | PS00649 | Group IV | SRR23629673 |
| 1 | rep2 | 3.08 | PS00649 | Group IV | SRR23629673 |
| 2 | rep2 | 3.08 | PS00649 | Group IV | SRR23629673 |
| 4 | rep2 | 3.21 | PS00649 | Group IV | SRR23629673 |
| 6 | rep2 | 3.31 | PS00649 | Group IV | SRR23629673 |
| 8 | rep2 | 3.36 | PS00649 | Group IV | SRR23629673 |
| 14 | rep2 | 4.12 | PS00649 | Group IV | SRR23629673 |
| 15 | rep2 | 4.22 | PS00649 | Group IV | SRR23629673 |
| 16 | rep2 | 4.39 | PS00649 | Group IV | SRR23629673 |
| 18 | rep2 | 4.95 | PS00649 | Group IV | SRR23629673 |
| 20 | rep2 | 5.55 | PS00649 | Group IV | SRR23629673 |
| 22 | rep2 | 6.14 | PS00649 | Group IV | SRR23629673 |
| 24 | rep2 | 6.41 | PS00649 | Group IV | SRR23629673 |
| 27 | rep2 | 6.53 | PS00649 | Group IV | SRR23629673 |
| 30 | rep2 | 6.81 | PS00649 | Group IV | SRR23629673 |

10°C growth data

| Time (h) | Rep | Log Count | Isolate | *panC* phylogenetic group | Accession Number |
|---|---|---|---|---|---|
| 0 | rep1 | 3.22 | PS00193 | Group II | SRR5185018 |
| 6 | rep1 | 3.22 | PS00193 | Group II | SRR5185018 |
| 24 | rep1 | 3.27 | PS00193 | Group II | SRR5185018 |
| 48 | rep1 | 3.30 | PS00193 | Group II | SRR5185018 |
| 96 | rep1 | 3.60 | PS00193 | Group II | SRR5185018 |
| 192 | rep1 | 4.05 | PS00193 | Group II | SRR5185018 |
| 240 | rep1 | 4.69 | PS00193 | Group II | SRR5185018 |
| 288 | rep1 | 6.05 | PS00193 | Group II | SRR5185018 |
| 384 | rep1 | 5.91 | PS00193 | Group II | SRR5185018 |
| 504 | rep1 | 5.71 | PS00193 | Group II | SRR5185018 |
| 0 | rep2 | 3.14 | PS00193 | Group II | SRR5185018 |
| 6 | rep2 | 3.14 | PS00193 | Group II | SRR5185018 |
| 24 | rep2 | 3.17 | PS00193 | Group II | SRR5185018 |
| 48 | rep2 | 3.25 | PS00193 | Group II | SRR5185018 |
| 96 | rep2 | 3.47 | PS00193 | Group II | SRR5185018 |
| 192 | rep2 | 4.03 | PS00193 | Group II | SRR5185018 |
| 240 | rep2 | 4.63 | PS00193 | Group II | SRR5185018 |
| 288 | rep2 | 6.07 | PS00193 | Group II | SRR5185018 |
| 384 | rep2 | 5.95 | PS00193 | Group II | SRR5185018 |
| 504 | rep2 | 5.59 | PS00193 | Group II | SRR5185018 |
| 0 | rep1 | 3.00 | PS00194 | Group VII | ASM225094v2 |
| 6 | rep1 | 3.06 | PS00194 | Group VII | ASM225094v2 |

| | | | | | |
|---|---|---|---|---|---|
| 24 | rep1 | 3.26 | PS00194 | Group VII | ASM225094v2 |
| 48 | rep1 | 3.34 | PS00194 | Group VII | ASM225094v2 |
| 96 | rep1 | 3.95 | PS00194 | Group VII | ASM225094v2 |
| 192 | rep1 | 6.46 | PS00194 | Group VII | ASM225094v2 |
| 240 | rep1 | 6.62 | PS00194 | Group VII | ASM225094v2 |
| 288 | rep1 | 6.65 | PS00194 | Group VII | ASM225094v2 |
| 384 | rep1 | 7.11 | PS00194 | Group VII | ASM225094v2 |
| 504 | rep1 | 6.42 | PS00194 | Group VII | ASM225094v2 |
| 0 | rep2 | 2.99 | PS00194 | Group VII | ASM225094v2 |
| 6 | rep2 | 3.03 | PS00194 | Group VII | ASM225094v2 |
| 24 | rep2 | 3.02 | PS00194 | Group VII | ASM225094v2 |
| 48 | rep2 | 3.03 | PS00194 | Group VII | ASM225094v2 |
| 96 | rep2 | 3.10 | PS00194 | Group VII | ASM225094v2 |
| 192 | rep2 | 6.43 | PS00194 | Group VII | ASM225094v2 |
| 240 | rep2 | 6.74 | PS00194 | Group VII | ASM225094v2 |
| 288 | rep2 | 6.79 | PS00194 | Group VII | ASM225094v2 |
| 384 | rep2 | 6.86 | PS00194 | Group VII | ASM225094v2 |
| 504 | rep2 | 6.28 | PS00194 | Group VII | ASM225094v2 |
| 0 | rep1 | 3.04 | PS00402 | Group IV | SRR23629691 |
| 6 | rep1 | 3.15 | PS00402 | Group IV | SRR23629691 |
| 24 | rep1 | 3.03 | PS00402 | Group IV | SRR23629691 |
| 48 | rep1 | 3.06 | PS00402 | Group IV | SRR23629691 |
| 96 | rep1 | 3.20 | PS00402 | Group IV | SRR23629691 |
| 192 | rep1 | 4.80 | PS00402 | Group IV | SRR23629691 |
| 240 | rep1 | 6.53 | PS00402 | Group IV | SRR23629691 |
| 288 | rep1 | 6.59 | PS00402 | Group IV | SRR23629691 |
| 384 | rep1 | 6.65 | PS00402 | Group IV | SRR23629691 |
| 504 | rep1 | 6.32 | PS00402 | Group IV | SRR23629691 |
| 0 | rep2 | 3.14 | PS00402 | Group IV | SRR23629691 |
| 6 | rep2 | 3.15 | PS00402 | Group IV | SRR23629691 |
| 24 | rep2 | 3.39 | PS00402 | Group IV | SRR23629691 |
| 48 | rep2 | 3.42 | PS00402 | Group IV | SRR23629691 |
| 96 | rep2 | 3.49 | PS00402 | Group IV | SRR23629691 |
| 192 | rep2 | 3.95 | PS00402 | Group IV | SRR23629691 |
| 240 | rep2 | 4.41 | PS00402 | Group IV | SRR23629691 |
| 288 | rep2 | 5.48 | PS00402 | Group IV | SRR23629691 |
| 384 | rep2 | 5.66 | PS00402 | Group IV | SRR23629691 |
| 504 | rep2 | 3.96 | PS00402 | Group IV | SRR23629691 |
| 0 | rep1 | 3.07 | PS00407 | Group IV | SRR13988985 |
| 6 | rep1 | 3.08 | PS00407 | Group IV | SRR13988985 |
| 24 | rep1 | 3.04 | PS00407 | Group IV | SRR13988985 |
| 48 | rep1 | 3.07 | PS00407 | Group IV | SRR13988985 |
| 96 | rep1 | 4.10 | PS00407 | Group IV | SRR13988985 |
| 192 | rep1 | 5.26 | PS00407 | Group IV | SRR13988985 |
| 240 | rep1 | 5.94 | PS00407 | Group IV | SRR13988985 |

| | | | | | |
|---|---|---|---|---|---|
| 288 | rep1 | 6.36 | PS00407 | Group IV | SRR13988985 |
| 384 | rep1 | 6.67 | PS00407 | Group IV | SRR13988985 |
| 504 | rep1 | 6.39 | PS00407 | Group IV | SRR13988985 |
| 0 | rep2 | 3.29 | PS00407 | Group IV | SRR13988985 |
| 6 | rep2 | 3.28 | PS00407 | Group IV | SRR13988985 |
| 24 | rep2 | 3.34 | PS00407 | Group IV | SRR13988985 |
| 48 | rep2 | 3.50 | PS00407 | Group IV | SRR13988985 |
| 96 | rep2 | 3.63 | PS00407 | Group IV | SRR13988985 |
| 192 | rep2 | 3.84 | PS00407 | Group IV | SRR13988985 |
| 240 | rep2 | 4.58 | PS00407 | Group IV | SRR13988985 |
| 288 | rep2 | 5.68 | PS00407 | Group IV | SRR13988985 |
| 384 | rep2 | 5.83 | PS00407 | Group IV | SRR13988985 |
| 504 | rep2 | 4.72 | PS00407 | Group IV | SRR13988985 |
| 0 | rep1 | 3.73 | PS00413 | Group IV | SRR13988908 |
| 6 | rep1 | 3.75 | PS00413 | Group IV | SRR13988908 |
| 24 | rep1 | 3.56 | PS00413 | Group IV | SRR13988908 |
| 48 | rep1 | 4.86 | PS00413 | Group IV | SRR13988908 |
| 96 | rep1 | 6.80 | PS00413 | Group IV | SRR13988908 |
| 192 | rep1 | 7.39 | PS00413 | Group IV | SRR13988908 |
| 240 | rep1 | 6.61 | PS00413 | Group IV | SRR13988908 |
| 288 | rep1 | 6.66 | PS00413 | Group IV | SRR13988908 |
| 384 | rep1 | 6.63 | PS00413 | Group IV | SRR13988908 |
| 0 | rep2 | 3.63 | PS00413 | Group IV | SRR13988908 |
| 6 | rep2 | 3.63 | PS00413 | Group IV | SRR13988908 |
| 24 | rep2 | 3.76 | PS00413 | Group IV | SRR13988908 |
| 48 | rep2 | 3.79 | PS00413 | Group IV | SRR13988908 |
| 96 | rep2 | 5.74 | PS00413 | Group IV | SRR13988908 |
| 192 | rep2 | 6.49 | PS00413 | Group IV | SRR13988908 |
| 240 | rep2 | 6.67 | PS00413 | Group IV | SRR13988908 |
| 288 | rep2 | 6.89 | PS00413 | Group IV | SRR13988908 |
| 384 | rep2 | 6.01 | PS00413 | Group IV | SRR13988908 |
| 0 | rep1 | 3.39 | PS00433 | Group IV | SRR13988883 |
| 6 | rep1 | 3.45 | PS00433 | Group IV | SRR13988883 |
| 24 | rep1 | 3.53 | PS00433 | Group IV | SRR13988883 |
| 48 | rep1 | 3.64 | PS00433 | Group IV | SRR13988883 |
| 96 | rep1 | 3.94 | PS00433 | Group IV | SRR13988883 |
| 192 | rep1 | 5.79 | PS00433 | Group IV | SRR13988883 |
| 240 | rep1 | 6.19 | PS00433 | Group IV | SRR13988883 |
| 288 | rep1 | 6.25 | PS00433 | Group IV | SRR13988883 |
| 384 | rep1 | 6.10 | PS00433 | Group IV | SRR13988883 |
| 0 | rep2 | 3.73 | PS00433 | Group IV | SRR13988883 |
| 6 | rep2 | 3.76 | PS00433 | Group IV | SRR13988883 |
| 24 | rep2 | 3.85 | PS00433 | Group IV | SRR13988883 |
| 48 | rep2 | 3.88 | PS00433 | Group IV | SRR13988883 |
| 96 | rep2 | 3.83 | PS00433 | Group IV | SRR13988883 |

| | | | | | |
|---|---|---|---|---|---|
| 192 | rep2 | 5.54 | PS00433 | Group IV | SRR13988883 |
| 240 | rep2 | 5.86 | PS00433 | Group IV | SRR13988883 |
| 288 | rep2 | 6.15 | PS00433 | Group IV | SRR13988883 |
| 384 | rep2 | 5.98 | PS00433 | Group IV | SRR13988883 |
| 0 | rep1 | 2.97 | PS00457 | Group II | SRR13988862 |
| 6 | rep1 | 3.03 | PS00457 | Group II | SRR13988862 |
| 24 | rep1 | 3.15 | PS00457 | Group II | SRR13988862 |
| 48 | rep1 | 3.24 | PS00457 | Group II | SRR13988862 |
| 96 | rep1 | 3.62 | PS00457 | Group II | SRR13988862 |
| 192 | rep1 | 4.16 | PS00457 | Group II | SRR13988862 |
| 240 | rep1 | 5.66 | PS00457 | Group II | SRR13988862 |
| 288 | rep1 | 5.85 | PS00457 | Group II | SRR13988862 |
| 384 | rep1 | 6.06 | PS00457 | Group II | SRR13988862 |
| 504 | rep1 | 6.02 | PS00457 | Group II | SRR13988862 |
| 0 | rep2 | 3.05 | PS00457 | Group II | SRR13988862 |
| 6 | rep2 | 3.03 | PS00457 | Group II | SRR13988862 |
| 24 | rep2 | 3.07 | PS00457 | Group II | SRR13988862 |
| 48 | rep2 | 3.15 | PS00457 | Group II | SRR13988862 |
| 96 | rep2 | 3.29 | PS00457 | Group II | SRR13988862 |
| 192 | rep2 | 5.67 | PS00457 | Group II | SRR13988862 |
| 240 | rep2 | 6.24 | PS00457 | Group II | SRR13988862 |
| 288 | rep2 | 6.37 | PS00457 | Group II | SRR13988862 |
| 384 | rep2 | 6.34 | PS00457 | Group II | SRR13988862 |
| 504 | rep2 | N/A | PS00457 | Group II | SRR13988862 |
| 0 | rep1 | 3.26 | PS00474 | Group III | SRR13988850 |
| 6 | rep1 | 3.18 | PS00474 | Group III | SRR13988850 |
| 24 | rep1 | 3.11 | PS00474 | Group III | SRR13988850 |
| 48 | rep1 | 3.17 | PS00474 | Group III | SRR13988850 |
| 96 | rep1 | 3.22 | PS00474 | Group III | SRR13988850 |
| 192 | rep1 | 5.31 | PS00474 | Group III | SRR13988850 |
| 240 | rep1 | 6.13 | PS00474 | Group III | SRR13988850 |
| 288 | rep1 | 6.47 | PS00474 | Group III | SRR13988850 |
| 384 | rep1 | 6.42 | PS00474 | Group III | SRR13988850 |
| 0 | rep2 | 3.68 | PS00474 | Group III | SRR13988850 |
| 6 | rep2 | 3.62 | PS00474 | Group III | SRR13988850 |
| 24 | rep2 | 3.60 | PS00474 | Group III | SRR13988850 |
| 48 | rep2 | 3.67 | PS00474 | Group III | SRR13988850 |
| 96 | rep2 | 3.71 | PS00474 | Group III | SRR13988850 |
| 192 | rep2 | 3.97 | PS00474 | Group III | SRR13988850 |
| 240 | rep2 | 4.59 | PS00474 | Group III | SRR13988850 |
| 288 | rep2 | 5.41 | PS00474 | Group III | SRR13988850 |
| 384 | rep2 | 4.65 | PS00474 | Group III | SRR13988850 |
| 0 | rep1 | 3.60 | PS00495 | Group IV | SRR13988829 |
| 6 | rep1 | 3.66 | PS00495 | Group IV | SRR13988829 |
| 24 | rep1 | 3.79 | PS00495 | Group IV | SRR13988829 |

| | | | | | |
|---|---|---|---|---|---|
| 48 | rep1 | 3.75 | PS00495 | Group IV | SRR13988829 |
| 96 | rep1 | 4.28 | PS00495 | Group IV | SRR13988829 |
| 192 | rep1 | 5.14 | PS00495 | Group IV | SRR13988829 |
| 240 | rep1 | 6.16 | PS00495 | Group IV | SRR13988829 |
| 288 | rep1 | 6.40 | PS00495 | Group IV | SRR13988829 |
| 384 | rep1 | 6.51 | PS00495 | Group IV | SRR13988829 |
| 0 | rep2 | 3.02 | PS00495 | Group IV | SRR13988829 |
| 6 | rep2 | 2.88 | PS00495 | Group IV | SRR13988829 |
| 24 | rep2 | 3.04 | PS00495 | Group IV | SRR13988829 |
| 48 | rep2 | 3.29 | PS00495 | Group IV | SRR13988829 |
| 96 | rep2 | 3.46 | PS00495 | Group IV | SRR13988829 |
| 192 | rep2 | 6.27 | PS00495 | Group IV | SRR13988829 |
| 240 | rep2 | 6.46 | PS00495 | Group IV | SRR13988829 |
| 288 | rep2 | 6.88 | PS00495 | Group IV | SRR13988829 |
| 384 | rep2 | 5.72 | PS00495 | Group IV | SRR13988829 |
| 0 | rep1 | 3.14 | PS00518 | Group II | SRR13989003 |
| 6 | rep1 | 3.12 | PS00518 | Group II | SRR13989003 |
| 24 | rep1 | 3.31 | PS00518 | Group II | SRR13989003 |
| 48 | rep1 | 3.27 | PS00518 | Group II | SRR13989003 |
| 96 | rep1 | 4.26 | PS00518 | Group II | SRR13989003 |
| 192 | rep1 | 6.92 | PS00518 | Group II | SRR13989003 |
| 240 | rep1 | 6.93 | PS00518 | Group II | SRR13989003 |
| 288 | rep1 | 7.00 | PS00518 | Group II | SRR13989003 |
| 384 | rep1 | 6.92 | PS00518 | Group II | SRR13989003 |
| 0 | rep2 | 3.14 | PS00518 | Group II | SRR13989003 |
| 6 | rep2 | 3.15 | PS00518 | Group II | SRR13989003 |
| 24 | rep2 | 3.08 | PS00518 | Group II | SRR13989003 |
| 48 | rep2 | 3.12 | PS00518 | Group II | SRR13989003 |
| 96 | rep2 | 3.79 | PS00518 | Group II | SRR13989003 |
| 192 | rep2 | 6.24 | PS00518 | Group II | SRR13989003 |
| 240 | rep2 | 6.56 | PS00518 | Group II | SRR13989003 |
| 288 | rep2 | 6.77 | PS00518 | Group II | SRR13989003 |
| 384 | rep2 | 6.43 | PS00518 | Group II | SRR13989003 |
| 0 | rep1 | 3.11 | PS00536 | Group VII | SRR13988990 |
| 6 | rep1 | 3.12 | PS00536 | Group VII | SRR13988990 |
| 24 | rep1 | 3.52 | PS00536 | Group VII | SRR13988990 |
| 48 | rep1 | 3.37 | PS00536 | Group VII | SRR13988990 |
| 96 | rep1 | 4.26 | PS00536 | Group VII | SRR13988990 |
| 192 | rep1 | 6.85 | PS00536 | Group VII | SRR13988990 |
| 240 | rep1 | 6.85 | PS00536 | Group VII | SRR13988990 |
| 288 | rep1 | 6.82 | PS00536 | Group VII | SRR13988990 |
| 384 | rep1 | 6.71 | PS00536 | Group VII | SRR13988990 |
| 0 | rep2 | 3.06 | PS00536 | Group VII | SRR13988990 |
| 6 | rep2 | 3.09 | PS00536 | Group VII | SRR13988990 |
| 24 | rep2 | 3.39 | PS00536 | Group VII | SRR13988990 |

| | | | | | |
|---|---|---|---|---|---|
| 48 | rep2 | 3.42 | PS00536 | Group VII | SRR13988990 |
| 96 | rep2 | 3.66 | PS00536 | Group VII | SRR13988990 |
| 192 | rep2 | 5.59 | PS00536 | Group VII | SRR13988990 |
| 240 | rep2 | 6.55 | PS00536 | Group VII | SRR13988990 |
| 288 | rep2 | 6.31 | PS00536 | Group VII | SRR13988990 |
| 384 | rep2 | 6.04 | PS00536 | Group VII | SRR13988990 |
| 0 | rep1 | 3.52 | PS00564 | Group II | SRR13988960 |
| 6 | rep1 | 3.13 | PS00564 | Group II | SRR13988960 |
| 24 | rep1 | 3.68 | PS00564 | Group II | SRR13988960 |
| 48 | rep1 | 4.07 | PS00564 | Group II | SRR13988960 |
| 96 | rep1 | 6.15 | PS00564 | Group II | SRR13988960 |
| 192 | rep1 | 6.67 | PS00564 | Group II | SRR13988960 |
| 240 | rep1 | 6.59 | PS00564 | Group II | SRR13988960 |
| 288 | rep1 | 6.85 | PS00564 | Group II | SRR13988960 |
| 384 | rep1 | 6.52 | PS00564 | Group II | SRR13988960 |
| 0 | rep2 | 3.65 | PS00564 | Group II | SRR13988960 |
| 6 | rep2 | 3.64 | PS00564 | Group II | SRR13988960 |
| 24 | rep2 | 3.68 | PS00564 | Group II | SRR13988960 |
| 48 | rep2 | 3.71 | PS00564 | Group II | SRR13988960 |
| 96 | rep2 | 5.66 | PS00564 | Group II | SRR13988960 |
| 192 | rep2 | 6.35 | PS00564 | Group II | SRR13988960 |
| 240 | rep2 | 6.52 | PS00564 | Group II | SRR13988960 |
| 288 | rep2 | 6.57 | PS00564 | Group II | SRR13988960 |
| 384 | rep2 | 6.54 | PS00564 | Group II | SRR13988960 |
| 0 | rep1 | 3.92 | PS00570 | Group V | SRR13988955 |
| 6 | rep1 | 3.93 | PS00570 | Group V | SRR13988955 |
| 24 | rep1 | 3.97 | PS00570 | Group V | SRR13988955 |
| 48 | rep1 | 4.37 | PS00570 | Group V | SRR13988955 |
| 96 | rep1 | 6.14 | PS00570 | Group V | SRR13988955 |
| 192 | rep1 | 6.56 | PS00570 | Group V | SRR13988955 |
| 240 | rep1 | 6.48 | PS00570 | Group V | SRR13988955 |
| 288 | rep1 | 6.58 | PS00570 | Group V | SRR13988955 |
| 384 | rep1 | 6.38 | PS00570 | Group V | SRR13988955 |
| 0 | rep2 | 3.58 | PS00570 | Group V | SRR13988955 |
| 6 | rep2 | 3.61 | PS00570 | Group V | SRR13988955 |
| 24 | rep2 | 3.65 | PS00570 | Group V | SRR13988955 |
| 48 | rep2 | 3.68 | PS00570 | Group V | SRR13988955 |
| 96 | rep2 | 4.90 | PS00570 | Group V | SRR13988955 |
| 192 | rep2 | 6.78 | PS00570 | Group V | SRR13988955 |
| 240 | rep2 | 6.81 | PS00570 | Group V | SRR13988955 |
| 288 | rep2 | 6.73 | PS00570 | Group V | SRR13988955 |
| 384 | rep2 | 6.54 | PS00570 | Group V | SRR13988955 |
| 0 | rep1 | 3.24 | PS00638 | Group V | SRR13988845 |
| 6 | rep1 | 3.26 | PS00638 | Group V | SRR13988845 |
| 24 | rep1 | 3.96 | PS00638 | Group V | SRR13988845 |

| | | | | | |
|---|---|---|---|---|---|
| 48 | rep1 | 4.01 | PS00638 | Group V | SRR13988845 |
| 96 | rep1 | 4.20 | PS00638 | Group V | SRR13988845 |
| 192 | rep1 | 5.94 | PS00638 | Group V | SRR13988845 |
| 240 | rep1 | 6.15 | PS00638 | Group V | SRR13988845 |
| 288 | rep1 | 6.43 | PS00638 | Group V | SRR13988845 |
| 384 | rep1 | 6.21 | PS00638 | Group V | SRR13988845 |
| 0 | rep2 | 3.60 | PS00638 | Group V | SRR13988845 |
| 6 | rep2 | 3.64 | PS00638 | Group V | SRR13988845 |
| 24 | rep2 | 3.60 | PS00638 | Group V | SRR13988845 |
| 48 | rep2 | 3.61 | PS00638 | Group V | SRR13988845 |
| 96 | rep2 | 3.75 | PS00638 | Group V | SRR13988845 |
| 192 | rep2 | 5.56 | PS00638 | Group V | SRR13988845 |
| 240 | rep2 | 5.86 | PS00638 | Group V | SRR13988845 |
| 288 | rep2 | 5.98 | PS00638 | Group V | SRR13988845 |
| 384 | rep2 | 4.90 | PS00638 | Group V | SRR13988845 |
| 0 | rep1 | 3.22 | PS00649 | Group IV | SRR23629673 |
| 6 | rep1 | 3.19 | PS00649 | Group IV | SRR23629673 |
| 24 | rep1 | 3.23 | PS00649 | Group IV | SRR23629673 |
| 48 | rep1 | 3.32 | PS00649 | Group IV | SRR23629673 |
| 96 | rep1 | 3.47 | PS00649 | Group IV | SRR23629673 |
| 192 | rep1 | 5.94 | PS00649 | Group IV | SRR23629673 |
| 240 | rep1 | 6.18 | PS00649 | Group IV | SRR23629673 |
| 288 | rep1 | 6.15 | PS00649 | Group IV | SRR23629673 |
| 384 | rep1 | 6.25 | PS00649 | Group IV | SRR23629673 |
| 504 | rep1 | 5.98 | PS00649 | Group IV | SRR23629673 |
| 0 | rep2 | 3.53 | PS00649 | Group IV | SRR23629673 |
| 6 | rep2 | 3.66 | PS00649 | Group IV | SRR23629673 |
| 24 | rep2 | 3.47 | PS00649 | Group IV | SRR23629673 |
| 48 | rep2 | 3.47 | PS00649 | Group IV | SRR23629673 |
| 96 | rep2 | 3.98 | PS00649 | Group IV | SRR23629673 |
| 192 | rep2 | 5.53 | PS00649 | Group IV | SRR23629673 |
| 240 | rep2 | 6.22 | PS00649 | Group IV | SRR23629673 |
| 288 | rep2 | 6.44 | PS00649 | Group IV | SRR23629673 |
| 384 | rep2 | 6.58 | PS00649 | Group IV | SRR23629673 |
| 504 | rep2 | 6.32 | PS00649 | Group IV | SRR23629673 |

14

| 16°C growth data | | | | | |
|---|---|---|---|---|---|
| Time (h) | Rep | Log Count | Isolate | *panC* phylogenetic group | Accession Number |
| 0 | rep1 | 3.87 | PS00125 | Group I | SRR23629669 |
| 1 | rep1 | 3.97 | PS00125 | Group I | SRR23629669 |
| 2 | rep1 | 4.00 | PS00125 | Group I | SRR23629669 |
| 4 | rep1 | 4.07 | PS00125 | Group I | SRR23629669 |

| | | | | | |
|---|---|---|---|---|---|
| 6 | rep1 | 4.14 | PS00125 | Group I | SRR23629669 |
| 8 | rep1 | 4.25 | PS00125 | Group I | SRR23629669 |
| 14 | rep1 | 4.33 | PS00125 | Group I | SRR23629669 |
| 18 | rep1 | 5.59 | PS00125 | Group I | SRR23629669 |
| 24 | rep1 | 5.88 | PS00125 | Group I | SRR23629669 |
| 48 | rep1 | 6.19 | PS00125 | Group I | SRR23629669 |
| 96 | rep1 | 6.42 | PS00125 | Group I | SRR23629669 |
| 0 | rep2 | 3.85 | PS00125 | Group I | SRR23629669 |
| 1 | rep2 | 3.87 | PS00125 | Group I | SRR23629669 |
| 2 | rep2 | 3.90 | PS00125 | Group I | SRR23629669 |
| 4 | rep2 | 4.02 | PS00125 | Group I | SRR23629669 |
| 6 | rep2 | 4.16 | PS00125 | Group I | SRR23629669 |
| 8 | rep2 | 4.20 | PS00125 | Group I | SRR23629669 |
| 14 | rep2 | 4.42 | PS00125 | Group I | SRR23629669 |
| 18 | rep2 | 5.56 | PS00125 | Group I | SRR23629669 |
| 24 | rep2 | 5.83 | PS00125 | Group I | SRR23629669 |
| 48 | rep2 | 6.18 | PS00125 | Group I | SRR23629669 |
| 96 | rep2 | 6.42 | PS00125 | Group I | SRR23629669 |
| 0 | rep1 | 3.56 | PS00135 | Group I | ASM16145v1 |
| 1 | rep1 | 3.59 | PS00135 | Group I | ASM16145v1 |
| 2 | rep1 | 3.62 | PS00135 | Group I | ASM16145v1 |
| 4 | rep1 | 3.77 | PS00135 | Group I | ASM16145v1 |
| 6 | rep1 | 3.87 | PS00135 | Group I | ASM16145v1 |
| 8 | rep1 | 3.91 | PS00135 | Group I | ASM16145v1 |
| 14 | rep1 | 4.23 | PS00135 | Group I | ASM16145v1 |
| 18 | rep1 | 5.59 | PS00135 | Group I | ASM16145v1 |
| 24 | rep1 | 5.95 | PS00135 | Group I | ASM16145v1 |
| 48 | rep1 | 6.30 | PS00135 | Group I | ASM16145v1 |
| 96 | rep1 | 6.44 | PS00135 | Group I | ASM16145v1 |
| 0 | rep2 | 3.63 | PS00135 | Group I | ASM16145v1 |
| 1 | rep2 | 3.64 | PS00135 | Group I | ASM16145v1 |
| 2 | rep2 | 3.68 | PS00135 | Group I | ASM16145v1 |
| 4 | rep2 | 3.73 | PS00135 | Group I | ASM16145v1 |
| 6 | rep2 | 3.82 | PS00135 | Group I | ASM16145v1 |
| 8 | rep2 | 3.94 | PS00135 | Group I | ASM16145v1 |
| 14 | rep2 | 4.39 | PS00135 | Group I | ASM16145v1 |
| 18 | rep2 | 5.72 | PS00135 | Group I | ASM16145v1 |
| 24 | rep2 | 6.01 | PS00135 | Group I | ASM16145v1 |
| 48 | rep2 | 6.17 | PS00135 | Group I | ASM16145v1 |
| 96 | rep2 | 6.36 | PS00135 | Group I | ASM16145v1 |
