## Supplement Figure 1 for "Exposure assessment suggests some cytotoxic *Bacillus cereus* group genotypes can grow over 3 logs in HTST milk throughout the shelf life at temperature abuse conditions"

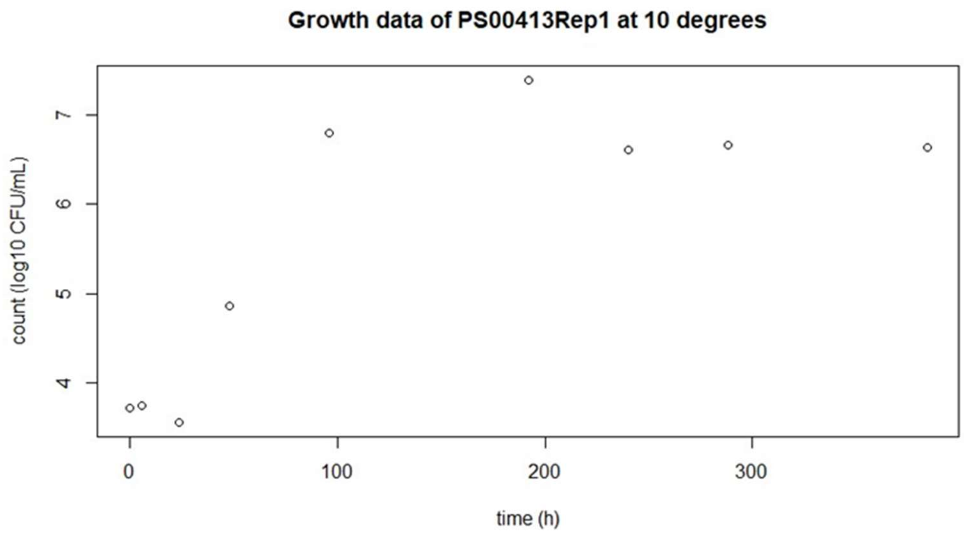


**Figure S1.** Plotted growth data of PS00413 (Rep1) at 10°C. The last three data points at 240, 288, 384 h indicate the death phase.
