## Supplementary figures and images for "Exposure assessment suggests some cytotoxic *Bacillus cereus* group genotypes can grow over 3 logs in HTST milk throughout the shelf life at temperature abuse conditions"

### Supplement Figure 2

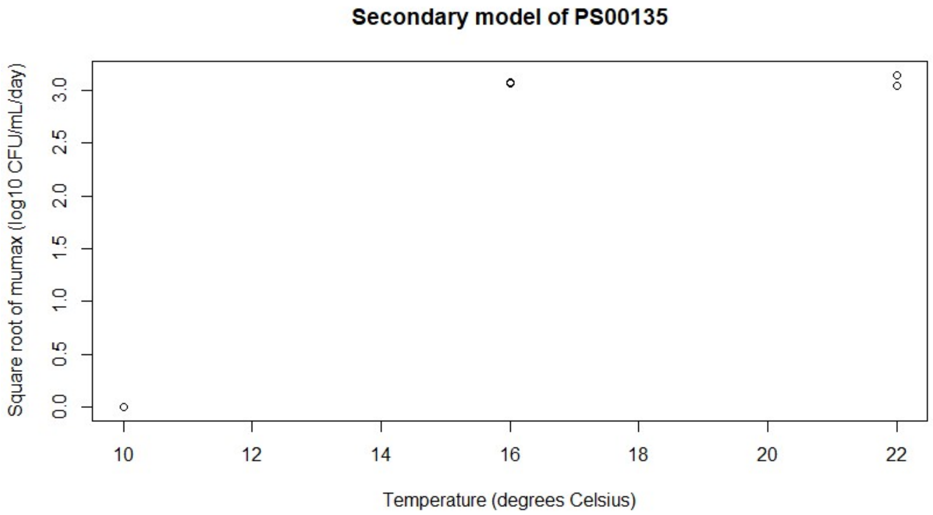


**Figure S2.** The secondary model of isolate PS00135 was non-linear.
